# Accurate Estimation of Molecular Counts from Amplicon Sequence Data with Unique Molecular Identifiers

**DOI:** 10.1101/2022.06.12.495839

**Authors:** Xiyu Peng, Karin S Dorman

**Affiliations:** Department of Epidemiology and Biostatistics, Memorial Sloan Kettering Cancer Center; Department of Statistics, Iowa State University, Ames, IA 50011, USA; Bioinformatics and Computational Biology Program, Iowa State University, Ames, IA 50011, USA; Department of Genetics, Development and Cell Biology, Iowa State University, Ames, IA 50011, USA

**Keywords:** unique molecular identifier, categorical data clustering, compositional data quantification, genetic variants, Hidden Markov Model, sparse parameter estimation

## Abstract

**Motivation:** Amplicon sequencing is widely applied to explore heterogeneity and rare variants in genetic populations. Resolving true biological variants and quantifying their abundance is crucial for downstream analyses, but measured abundances are distorted by stochasticity and bias in amplification, plus errors during Polymerase Chain Reaction (PCR) and sequencing. One solution attaches Unique Molecular Identifiers (UMIs) to sample sequences before amplification eliminating amplification bias by clustering reads on UMI and counting clusters to quantify abundance. While modern methods improve over naïve clustering by UMI identity, most do not account for UMI reuse, or collision, and they do not adequately model PCR and sequencing errors in the UMIs and sample sequences.

**Results:** We introduce Deduplication and accurate Abundance estimation with UMIs (DAUMI), a probabilistic framework to detect true biological sequences and accurately estimate their deduplicated abundance from amplicon sequence data. DAUMI recognizes UMI collision, even on highly similar sequences, and detects and corrects most PCR and sequencing errors in the UMI and sampled sequences. DAUMI performs better on simulated and real data compared to other UMI-aware clustering methods.

**Availability:** Source code is available at https://github.com/xiyupeng/AmpliCI-UMI.

## 1 Introduction

Amplicon sequencing has been widely applied to explore complex genetic populations, such as viral quasispecies [66, 47, 22, 3, 8], microbial communities [17, 24, 6], cancer tumors [36, 49], immune receptor repertoires [50, 62, 44], and engineered barcodes [4, 26, 33]. These experiments generate very high coverage sequence data for targeted genomic regions, enabling detection of variants with single nucleotide differences or low abundance [5, 18]. However, to attach sequencing adapters [43] and achieve deep coverage, the targeted regions are amplified by Polymerase Chain Reaction (PCR) and then sequenced using error-prone high throughput sequencing technologies. Stochasticity and bias in amplification along with errors generated during PCR and sequencing may obscure the true variants and distort measured abundances [25, 63]. Bias is especially problematic in low-biomass samples, like single cell samples, where relatively few template molecules are intensely amplified through many PCR cycles [60, 29]. Errors, chimeras and amplification bias accelerate with more PCR cycles [60, 63], burying real sequences in increasing levels of sequence noise.

The solution to PCR bias is to identify and remove PCR-duplicated molecules, but it is not easy to achieve, especially for amplicon data. The common strategy to assume duplicate reads are those mapping to the same reference location [32, 12, 16] may introduce more bias than it removes [30], and is not possible for amplicons. Attaching Unique Molecular Identifiers (UMIs) to sampled molecules before amplification was suggested [21] and implemented for Sanger sequencing [35], then high throughput sequencing [31, 28, 29]. If UMI tags are unique and unadulterated by library preparation and sequencing, then reads with the same UMI tag are amplified products of the same molecule. UMIs lower the limit of detection for ultra-low frequency variants [28, 64], eliminate amplification bias [29, 22], and UMI counts (deduplicated abundances) are cleaner than raw read count data and simplify downstream statistical analysis [61, 27, 59].

Early processing pipelines [62, 51, 56, 9] clustered reads by UMI identity and generated a consensus sample sequence per cluster. Unfortunately, UMIs can *collide*, identically marking two or more sampled molecules [10] and leading to undercount, loss or misestimation of sampled variants. And UMIs experience both PCR and sequencing errors, leading to discovery of false (error) variants and inflating the estimated number of sampled molecules [52]. Modern UMI-aware deduplication or quantification tools account for errors in UMIs, and sometimes, UMI collisions. Calib [37] clusters whole reads, the UMI and sample sequence, by sequence similarity. Use of whole reads can resolve UMI collisions attached to distinct sample sequences, but distance-based clustering without accounting for abundance of similar sequences is difficult to callibrate, likely to over-merge clusters if reads are similar or under-merge multi-error misreads of abundant haplotypes [6, 1, 38]. Starcode-umi [68] clusters UMIs and sample sequences separately, applying thresholds to distance and cluster abundances that need careful tuning [37], then reconstructs error-free reads from denoised UMIs and sample sequences. UMI-tools [52] clusters UMIs on a network, accounting for both abundance and similarity. Intended for single cell RNA-seq, it avoids UMI collision by partitioning UMIs on reference mapping position, but no such partition is possible for amplicon data.

We introduce DAUMI, Deduplication and accurate Abundance estimation with Unique Molecular Identifiers, a novel probabilistic framework to detect true biological sequences and accurately estimate their deduplicated abundance from amplicon sequence data. DAUMI detects and corrects PCR and sequencing errors (Figure 1a) better than traditional consensus methods. It also recognizes UMI collision, even on highly similar sample sequences (Supplementary Figure S1). On simulated and real data, DAUMI identifies fewer false positives and achieves similar or better abundance estimation than current approaches, particularly achieving higher accuracy on datasets with UMI collision.

**Fig. 1:**
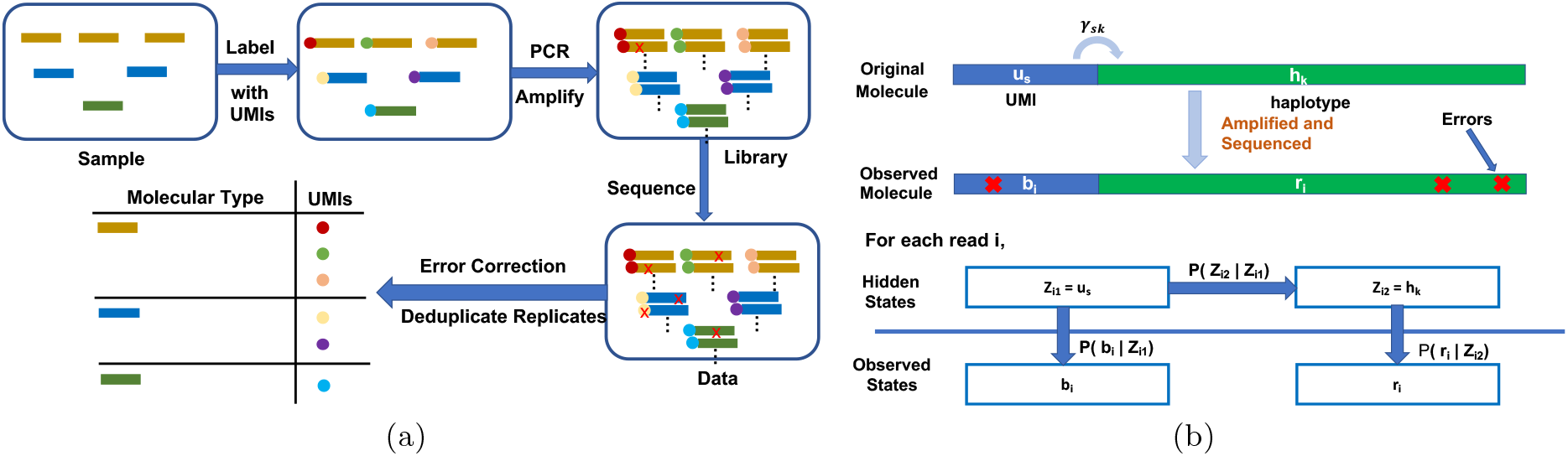
(a) Deduplication and error correction with UMIs. Red crosses are errors generated by PCR or sequencing. Our goal is to resolve the true biological sequences in the sample, as well as their sampled absolute abundance. (b) A one-step Hidden Markov Model (HMM). Each read has two hidden states, true UMI ***u***_*s*_ and true haplotype sequence ***h***_*k*_, and two observed states, observed UMI ***b***_*i*_ and observed sample sequence ***r***_*i*_. Both ***b***_*i*_ and ***r***_*i*_ are observations of ***u***_*s*_ and ***h***_*k*_ with error. Transition matrix ***Γ*** models the attachment of true UMI sequence ***u***_*s*_ to true haplotype sequence ***h***_*k*_.

## 2 Methods

We start with a dataset of *n* single-end reads, where the *i*th read contains an observed UMI ***b***_*i*_ and sample sequence ***r***_*i*_. A pair ***Z***_*i*_ = (*Z*_*i*1_, *Z*_*i*2_) of hidden variables for read *i* encode the unknown source of UMI sequence ***b***_*i*_, drawn from a set of *N* true UMIs, 𝒰 = {***u***_1_, ***u***_2_, …, ***u***_*N*_}, and the unknown source of sample sequence ***r***_*i*_, drawn from a set of *K* true sample sequences (haplotypes), ℋ = {***h***_1_, ***h***_2_, …, ***h***_*K*_}.

### 2.1 Model

Each read *i* is assumed to be an independent realization of a one-step Hidden Markov Model (Figure 1b). For ***b*** = (***b***_1_, ***b***_2_, …, ***b***_*n*_) and ***r*** = (***r***_1_, ***r***_2_, …, ***r***_*n*_), the observed log likelihood *l*(***θ*** | ***r, b***) is

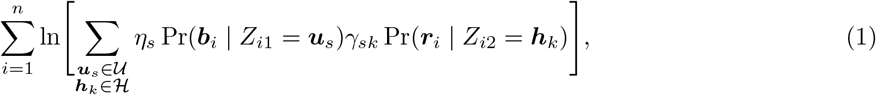

where ***θ*** = {***η, Γ***, ℋ, 𝒰} are model parameters. Parameters ***η*** = (*η*_1_, *η*_2_, …, *η*_*N*_) are the mixing proportions over the UMIs, which vary because of stochastic amplification, differences in initial UMI abundances by collision, and inclusion of low-frequency error UMIs in 𝒰. Matrix ***Γ*** = {*γ*_*sk*_}_*N*×*K*_ is a transition matrix used to model the attachment of true UMIs to haplotypes. For each unique UMI ***u***_*s*_, we expect 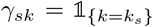, *i*.*e*., 0 unless *k* is *k*_*s*_, the index of the haplotype uniquely marked by UMI ***u***_*s*_. However, recombination during PCR (*e*.*g*., chimeras) [25, 41] and UMI collision, especially when UMIs are short [10] or non-uniformly distributed [39], reduce the sparsity of *Γ* by adding multiple positive transition probabilities *γ*_*sk*_ *>* 0 for the *s*th UMI.

In a well-executed experiment, however, transition matrix ***Γ*** will be sparse. To encourage sparsity, we maximize the observed *penalized* log likelihood, where penalty [7, 2],

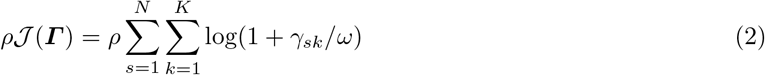

is subtracted from log likelihood (1). Parameters *ω* and *ρ* are positive constants the user must select. As *ω* → 0, the penalty becomes 𝓁_0_-like, such that transitions without support in the data are driven to 0. With *ω* small, the penalty induces sparseness in ***Γ*** by setting 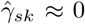 when the expected amplified count of molecule (***u***_*s*_, ***h***_*k*_) drops below threshold *ρ*. UMI collisions are evident when expected counts of 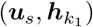 *and* 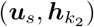 exceed *ρ*. Because both sequence similarity and abundance determine the expected counts, DAUMI can identify UMI collisions even on highly similar sampled molecules (Supplementary Figure S1). Further constraints are imposed on ***Γ*** to reduce computational complexity (see §2.2).

For emission probabilities Pr(***b***_*i*_ |*Z*_*i*1_) and Pr(***r***_*i*_ |*Z*_*i*2_), we use an existing model for sequencing errors [38]. Conditional on an alignment, the emission of observed sequence ***X*** from true sequence ***Y*** is

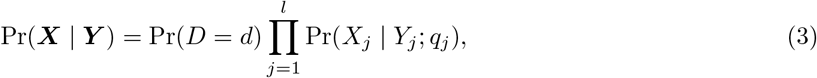

where *l* is the alignment length, *d* is the number of observed insertion/deletion (indel) events, and Pr(*X*_*j*_ | *Y*_*j*_; *q*_*j*_) is the probability (1 for indels) of generating nucleotide *X*_*j*_ from *Y*_*j*_ with quality score *q*_*j*_ at alignment position *j*. Indels follow a truncated Poisson(*δl*) with indel error rate *δ* per position per read, but we assume *d* = 0 in UMIs since indel errors are rare in these short, typically high quality parts of the reads. The goal of DAUMI is to estimate the true haplotypes ℋ and their abundance. We liberally initialize the sets ℋ and 𝒰of true haplotypes and UMIs as described in §2.2. Then, we iteratively maximize the penalized log likelihood to estimate model parameters using an expectation-maximization (EM) algorithm (see Supplementary Material §S1). ℋ and 𝒰 are fixed after initialization, but false UMIs and false haplotypes can be recognized and eliminated. Specifically after convergence, we discard UMIs with expected abundance 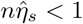 as likely error UMIs. Then, we estimate the deduplicated abundance of haplotype ***h***_*k*_ as 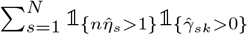.

### 2.2 Implementation

DAUMI is implemented in C. We describe EM algorithm initialization, approximations used for speed, and selection of penalty parameters.

#### Initialization

With 𝒰 and ℋgiven, we initialize mixing proportions ***η*** to the relative observed abundance of UMIs and transition matrix ***Γ*** to the observed count of each transition, normalizing row sums to one. To include all true UMIs and haplotypes in candidate pools 𝒰and ℋ, while excluding most error UMIs and haplotypes, we extend AmpliCI [38].

We set 𝒰to the “denoised” UMIs obtained by running AmpliCI on the UMI sequences (with option `−−umi`) while assuming quality scores provide PHRED error probabilities [15]. Though PHRED error probabilities are generally accurate [65], they slightly underestimate error rates [46, 51] and previous experience [6, 38] suggests this approach will liberally identify UMIs.

To populate ℋ, AmpliCI is applied to full reads (with option `−−trim_umi_length` to set UMI length), using an error profile estimated from (corrected) UMI clusters further split by a UMI-tools-like algorithm [52]. Specifically, if any unique sequence in a cluster achieves more than half the abundance of the most abundant sequence, we consider it a possible UMI collision and partition the cluster. The two sequences are promoted to new cluster centers, and each read in the original cluster is assigned to the closer center, by Hamming distance, or the more abundant center if equidistant. Also, any surviving error UMIs will tend to exist as a small cluster, where the true sequence and hence true errors may not be obvious, so we also exclude small clusters with total abundance below a threshold (default 2) in error estimation. Finally, the unique, “denoised” sequences, without UMI tags, form ℋ.

#### Approximations

For efficiency on large datasets, we truncate very small probabilities to zero to reduce computation and storage. Since Illumina error rates are low, *<*0.1% per nucleotide [57], we assume each true combination (***u***_*s*_, ***h***_*k*_) has been observed without error at least once and otherwise force *γ*_*sk*_ ≡ 0. Time and memory complexity increase with large numbers of surviving combinations of candidate haplotypes and UMIs, but most posterior probabilities *e*_*isk*_ are exceptionally small. In every E step, we only keep the top *T* (default 10) most likely candidates and renormalize to one for each read *i*, so

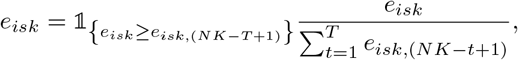

where *e*_*isk*,(*NK*−*T+*1)_ is the *T* th largest value among a *NK* entries. With these approximations, estimate 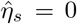 is possible, in which case we assume ***u***_*s*_ is an error UMI and the *s*th row of ***Γ*** is undefined and excluded.

#### Selection of the Penalty Parameters

Fixing *ω* = 10^−20^, for an 𝓁_0_ penalty, *ρ* is chosen to separate true from error UMIs. To automate *ρ* selection, we propose to model the observed UMI abundance distribution. Specifically, we modify the Galton-Watson branching process model [58] to account for amplification and sequencing error, and fit the model to the observed abundance of each unique UMI by minimizing the Kolmogorov–Smirnov statistic [54]. To accommodate long right tails caused by UMI collisions or other artefacts, we fit a right-truncated model. This solution works well for all datasets, except HIV V3, for which we manually choose *ρ* = 20 based on visual assessment. Details and limitations of this procedure are provided in Supplementary Material §S2. Supplementary Table S1 shows fitted models and selected *ρ* in all datasets except HIV V3.

### 2.3 Data Simulation

Figure 2 shows the simulation pipeline. Briefly, the relative abundance ***π*** of *K* = 25 haplotypes were obtained from power law distribution *f* (*x*) ∝ 0.5*x*^−0.5^𝕝_{0≤*x*≤1}_. Then *N* = 400 molecules were sampled according to a Multinomial(*N*, ***π***) distribution. Then, *N* distinct UMIs of nine nucleotides were randomly selected and attached to each molecule. For datasets with UMI collision, we resampled the *N* distinct UMIs *with replacement* before attaching them to the sampled molecules. The true sequences of the 400 UMIs and 25 haplotypes were randomly selected from the pools of UMIs and haplotypes found in an HIV V1-V2 dataset (Accession SRR2241783) [3] after AmpliCI denoising [38]. Next, tagged molecules were PCR amplified with PCR error probability per base, per cycle provided in [42] (10^−4^ – 10^−6^, depending on substitution). Simulation settings, including PCR efficiencies and cycle numbers, are provided in Table 1. ART Illumina [19] simulated single-end reads with fixed length 250nt, using error profile MSv1 to mimic Miseq sequences. Insertion and deletion rates were set to 2 × 10^−5^ per position.

**Table 1:** Simulated Datasets. *σ*: no. of PCR cycles; *χ*: PCR efficiency; Mean: expected amplified abundance (1 + *χ*)^*σ*^; Var.: variance in amplified abundance (1 −*χ*)(1 + *χ*)^*σ*^((1 + *χ*)^*σ*^ −1)*/*(1 + *χ*) [40]; Hap.: no. (of 25) true haplotypes sampled, with no. of unsampled haplotypes (true abundance 0) in parenthesis; UMI: no. true unique UMIs; Mol.: no. unique sampled molecules; *ρ*: selected penalty parameter.

| Simulation | $\sigma$ | $\chi$ | Mean | Var. | Hap. | UMI | Mol. | $\rho$ |
| --- | --- | --- | --- | --- | --- | --- | --- | --- |
| 1 | 10 | 0.5 | 57.67 | 1089.20 | 22 (3) | 400 | 400 | 9 |
| 2 | 7 | 0.9 | 89.39 | 415.83 | 23 (2) | 400 | 400 | 45 |
| 3 | 7 | 0.9 | 89.39 | 415.83 | 22 (3) | 263 | 400 | 49 |
| 4 | 10 | 0.5 | 57.67 | 1089.20 | 22 (3) | 257 | 400 | 18 |

**Fig. 2:**
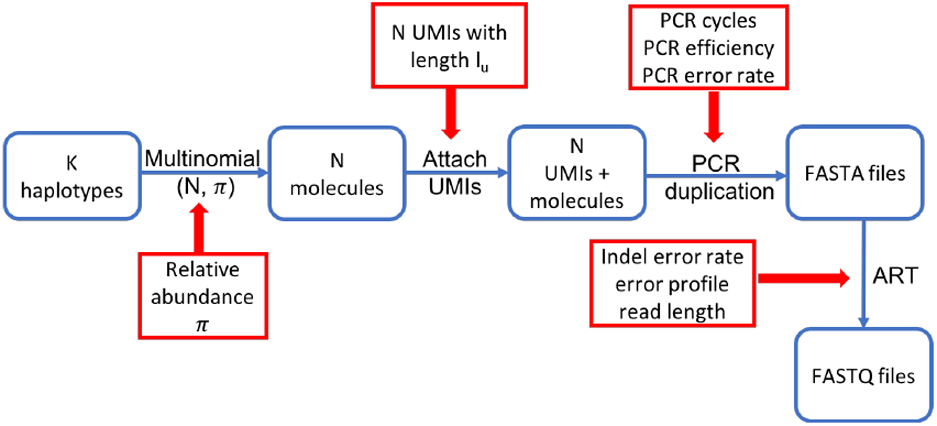
Data simulation pipeline. Total *N* molecules are tagged with UMIs, after being sampled from *K* haplotypes under a multinomial distribution (*N, π*). Then tagged molecular are amplified with a given efficiency for a given number of PCR cycles to mimic PCR amplification process. Lastly, ART is applied to simulate single-end reads to generate final FASTQ files.

### 2.4 Running Competing Methods

We run Calib [37], UMI-tools [52], Starcode-umi [68] and a “Naïve method” that clusters reads by UMI identity [62, 10] with default parameter settings, except as discussed here. Since Calib (v0.3.4) requires paired-end reads, we passed in the UMI-tagged reads as forward reads and the UMI-detagged reads as reverse reads (a simple hack suggested by the author). We disabled sequence trimming (`−−seq-trim 0`) in Starcode-umi (v1.3) to ensure a fair comparison with other algorithms, and set the maximum edit distance for UMI clustering as one (`−−umi-d 1`), a common setting for UMI clustering, instead of the default zero. For UMI-tools, we used the core algorithm from its Application Programming Interface (API), which is stable across versions, to cluster UMIs. For the Naïve method, sequences were clustered by UMI identity and singleton UMIs and their attached sample sequences were discarded as likely errors. For Calib, UMI-tools, and the Naïve method, we used the majority rule consensus sequence from each cluster to estimate the true UMI and haplotype (first seen of {`A,T,C,G}` when tied). For Starcode-umi, we used the canonical UMIs and sample sequences estimated during cluster formation. Supplementary Table S2 verifies chosen parameters were optimal or near-optimal for all methods (details in Supplementary Material §S3).

## 3 Results

We compared DAUMI to four UMI clustering methods, Calib [37], UMI-tools [52], Starcode-umi [68], and the Naïve method, on both simulated and real datasets.

### 3.1 Benchmarking on Simulated Data

To compare the methods, we simulated four datasets (Table 1). Of 25 true haplotypes, only 23 were sampled in Simulation 2 and 22 in the other simulations. Calib was run on subsets of Simulation 2 (50%), 3 (45%) and 4 (50%), since it failed to run on the whole dataset (true abundances have been adjusted accordingly).

DAUMI has superior performance in all four simulations (Figure 3 and Table 2). *Without* UMI collision, DAUMI, Calib, UMI-tools, and Naïve identify all true haplotypes, but DAUMI introduces fewer false positives (Figure 3, Simulations 1–2). Starcode-umi finds less true haplotypes and is the only method to overestimate abundance; its estimates also contain more noise. DAUMI excels on data *with* UMI collision (Figure 3, Simulations 3–4). UMI-tools and Naïve, which cannot detect collision, underestimate haplotype abundance, but Calib also underestimates abundance, possibly because it merges some UMI clusters with similar tagged reads.

**Table 2:**
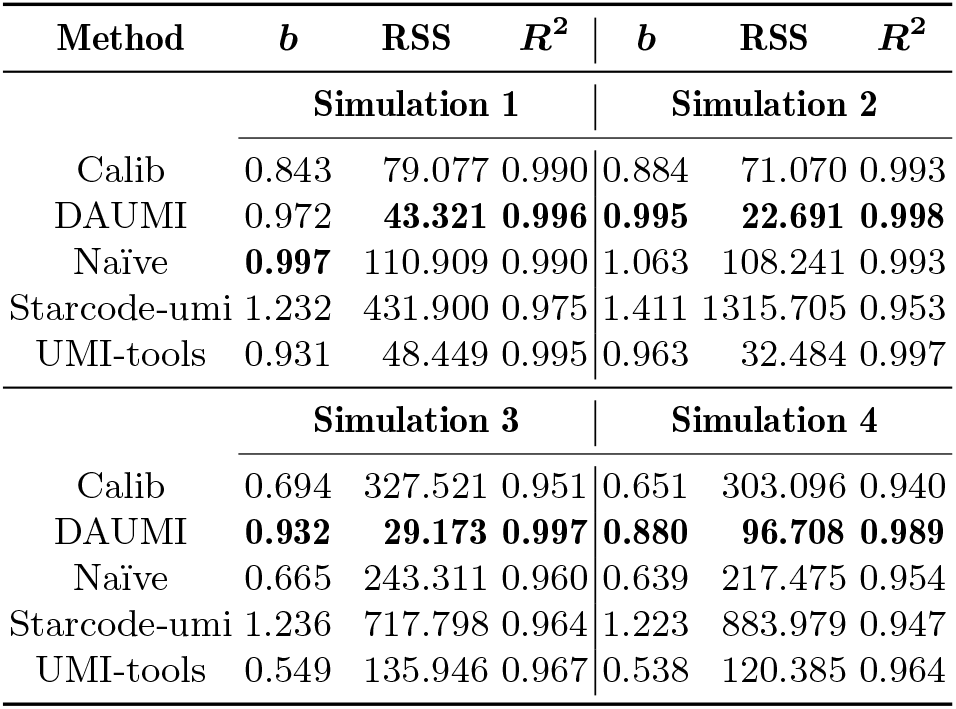
Abundance estimation for simulated data. From linear model Deduplicated Abundance = *b* × True Abundance fit, we report *b*: coefficient, *>* 1 for overestimation, *<* 1 for underestimation; RSS: residual sum of squares, 0 for perfect estimation; *R*^2^: proportion of variance explained, 1 for perfect estimation. The best achieved metric (column) is bolded per simulation.

**Fig. 3:**
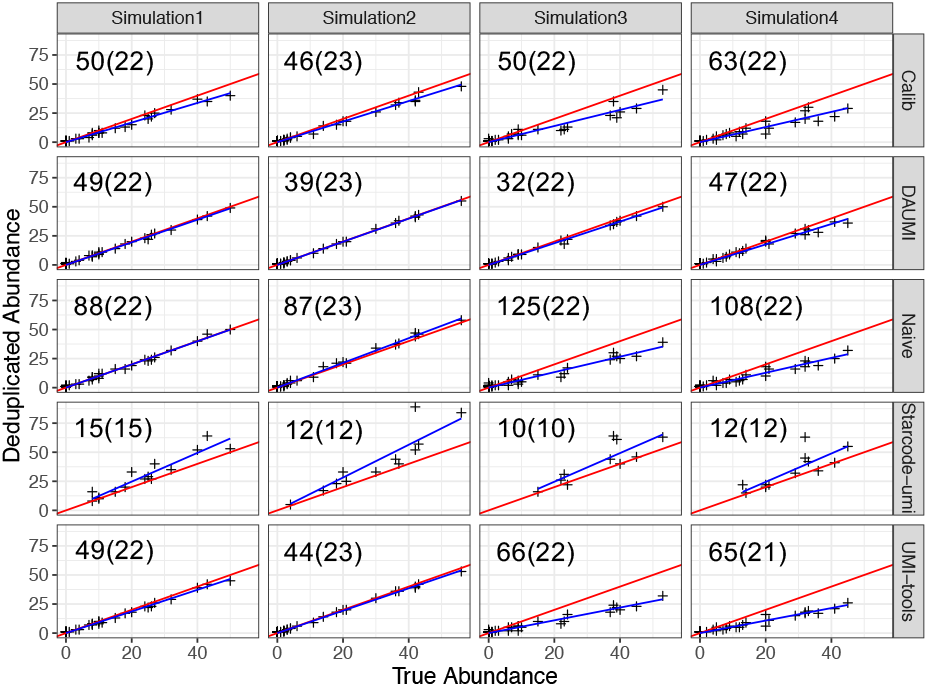
Abundance estimation on simulated data *without* (Simulations 1–2) or *with* (Simulations 3–4) UMI collision. Blue lines are fitted (*y* = *bx*) and red lines are expected (*y* = *x*). Number of estimated (true) haplotypes within each scatter plot. Estimates and goodness-of-fit in Table 2.

### 3.2 Application to Real Data

Only reverse reads, with attached UMIs, were analyzed. For V1V2, UMIs were extracted from the middle of raw reads using a custom method (Supplementary Material §S5). For V3, UMIs were extracted from the beginning of reverse reads, discarding the following adapter sequences. Finally, reads containing ambiguous nucleotides (1%) were removed, and all reads were trimmed to the same length. Detailed information on the datasets is in Supplementary Table S4.

To assess performance when the true haplotypes and abundances are unknown, we evaluated method consistency across a random split of each dataset into two equal-sized subsets (repeated five times). In Table 3 we report the total number of haplotypes found in each subset and two similarity metrics: Jaccard Index is a measure of agreement in found haplotypes [23]; Ruzicka Similarity [13] is further weighted by estimated abundances.

**Table 3:** Agreement across random partition of V1V2 and V3 datasets. The mean and standard deviation (in parentheses) of five replicates of: Hap1 and Hap2: no. inferred haplotypes in each subset (rounded to integer); Jaccard: Jaccard Index of inferred haplotype sets; Ruzicka: Ruzicka Similarity, abundance-weighted form of Jaccard Index. For DAUMI, *ρ* = 10 for V3, *ρ* = 7 for V1V2, both subsets. Best performance bolded per dataset.

| Dataset | Methods | Deduplicated abundance $\geq 1$ | | | | | | | | Deduplicated abundance $\geq 2$ | | | | | | | |
| --- | --- | --- | --- | --- | --- | --- | --- | --- | --- | --- | --- | --- | --- | --- | --- | --- | --- |
|  |  | Hap1 |  | Hap2 |  | Jaccard |  | Ruzicka |  | Hap1 |  | Hap2 |  | Jaccard |  | Ruzicka |  |
| V1V2 | Calib | 808 | (21) | 797 | (20) | 0.08 | (0.00) | 0.23 | (0.00) | 43 | (2) | 43 | (5) | 0.67 | (0.03) | 0.82 | (0.02) |
|  | DAUMI | 90 | (5) | 90 | (4) | **0.72** | (0.03) | **0.84** | (0.02) | 37 | (1) | 37 | (3) | **0.77** | (0.05) | 0.87 | (0.02) |
|  | Naïve | 293 | (10) | 303 | (7) | 0.29 | (0.01) | 0.57 | (0.01) | 43 | (2) | 43 | (3) | 0.75 | (0.03) | **0.88** | (0.03) |
|  | Starcode-umi | 2304 | (39) | 2311 | (55) | 0.003 | (0.000) | 0.19 | (0.00) | 15 | (1) | 13 | (1) | 0.73 | (0.05) | 0.80 | (0.02) |
|  | UMI-tools | 542 | (14) | 538 | (17) | 0.14 | (0.00) | 0.36 | (0.00) | 42 | (2) | 42 | (3) | **0.77** | (0.04) | 0.87 | (0.01) |
| V3 | Calib | 5048 | (42) | 5021 | (52) | 0.06 | (0.00) | 0.11 | (0.00) | 64 | (5) | 66 | (3) | 0.55 | (0.04) | 0.71 | (0.02) |
|  | DAUMI | 180 | (9) | 171 | (4) | **0.39** | (0.01) | **0.72** | (0.01) | 68 | (4) | 72 | (3) | **0.72** | (0.03) | **0.83** | (0.01) |
|  | Naïve | 1561 | (9) | 1543 | (10) | 0.06 | (0.00) | 0.22 | (0.00) | 91 | (4) | 84 | (3) | 0.64 | (0.04) | 0.74 | (0.02) |
|  | Starcode-umi | 15077 | (81) | 15097 | (76) | 0.02 | (0.00) | 0.09 | (0.00) | 96 | (6) | 103 | (4) | 0.08 | (0.01) | 0.77 | (0.02) |
|  | UMI-tools | 3718 | (37) | 3724 | (36) | 0.06 | (0.00) | 0.14 | (0.00) | 72 | (2) | 74 | (3) | 0.64 | (0.03) | 0.75 | (0.01) |

V1V2 amplifies the V1V2 region of the HIV-1 envelope (*env*) gene from an HIV superinfected individual who developed broadly neutralizing antibodies [3]. DAUMI achieves the highest performance and is competitive with UMI-tools and Naïve on the subset of haplotypes attached to at least two UMIs (Table 3). Though UMI collision is rare in these data, DAUMI and Starcode-umi detected a collision on two haplotypes with nine mismatches (Figure 4). Within this UMI cluster (126 sequences), DAUMI infers two haplotype sequences with high observed abundance (15 and 7) that also dominate other UMI clusters. Starcode-umi finds one of these haplotypes, but estimates the second to be a sequence never observed in the read data.

**Fig. 4:**
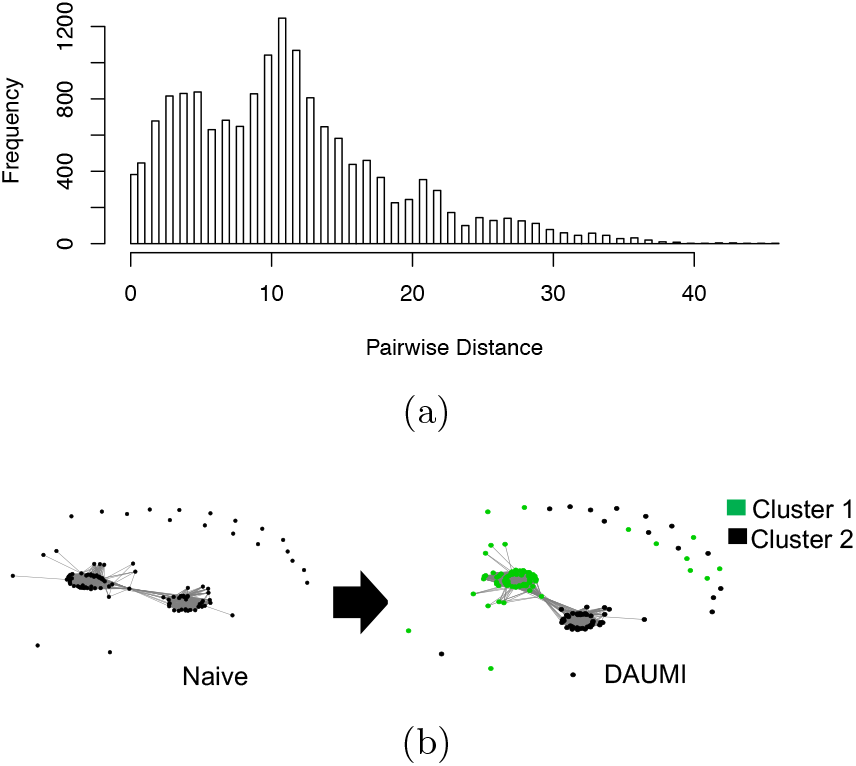
UMI collision in V1V2 data [3] resolved by DAUMI. (a) Distribution of pairwise distances between unique sequences labeled with UMI GTGTCGGTA, indicating at least two clusters linked this UMI. (b) DAUMI finds two clusters (green and black). Network nodes are unique sample sequences; edges link sequences within edit distance six.

V3 contains reads amplified from the V3 loop of *env* sampled from an HIV-1 infected patient undergoing immunotherapy with broadly neutralizing antibody 10-1074 [8]. DAUMI has the highest agreement in recovered haplotypes and estimated abundances. The larger performance advantage for DAUMI in V3 vs. V1V2 may result from a higher UMI collision rate in V3 data. Supplementary Figures S4–S5 show a much longer right tail despite lower PCR amplification, and Supplementary Figure S7 finds excess nucleotide T, suggesting degeneracy in the random UMIs.

All algorithms but Starcode-umi have higher agreement in recovered haplotypes and estimated abundances on V1V2 than V3 (Supplementary Figure S6 and S8). Though each algorithm identifies several haplotypes unique to that algorithm, estimated haplotypes with abundance no less than 2 are largely overlapped in V1V2. On both V1V2 and V3 datasets, Starcode-umi results are quite different from the other four algorithms, but it occassionally achieves high metrics, largely because it estimates a few haplotypes with very inflated abundances (Supplementary Figure S6).

## 4 Discussion

UMIs resolve sampled RNA or DNA molecules that are similar even when PCR and sequencing errors inflate or bias molecular counts. We have introduced DAUMI, a novel probabilistic framework to correct errors and deduplicate reads for accurate molecular counts of amplicon sequence data with UMIs. DAUMI correctly resolves the origin of more molecules than competing methods even with UMI collisions. Here, we discuss advantages, limitations, and directions for future research.

UMI collision happens, especially with short UMIs [10]. Low nucleotide diversity in high frequency UMIs was reported for single cell RNA-seq (scRNA-seq) [39], and we observed similar patterns in the V3 data. Most UMI-based quantification methods for scRNA-seq, *e*.*g*. UMI-tools derivative Alevin [53], map reads to avoid UMI collision, but mapping cannot resolve amplicons. Even in scRNA-seq, mapping may under-collapse clusters [53], since reads of the same molecule can map to slightly different genome positions [48]. Calib and Starcode-umi can detect UMI collision but were not competitive with DAUMI.

DAUMI and all methods compared here cannot detect errors introduced before UMI tagging, *e*.*g*. during cDNA synthesis. Downstream methods may detect such errors, like UMI-based variant callers [64, 51], which compare variation to background error rates after UMI clustering. Methods also struggle to detect early-cycle PCR errors, which are highly amplified and difficult to distinguish from true variation. When such errors occur in the UMI, methods may overestimate abundance by treating both UMIs as valid. Post hoc detection is possible by checking for similar UMIs attached to identical haplotypes. More likely, errors strike the sampled molecule. Other methods vary in their resolution of such cases, but DAUMI will treat the result as a UMI collision and overestimate the abundance of a false haplotype. If good experimental design has eliminated UMI collision, DAUMI abundance estimation could be altered to assign each UMI to one and only one haplotype, namely arg 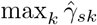 (with option `−−ncollision`), but when collision and early-cycle PCR error coexist, explicit modeling of the PCR process may be necessary.

DAUMI is moderately affected by the penalty parameter *ρ*, which is interpretable as a molecular count distinguishing error and true molecules. When there are UMI collisions, high *ρ* may merge similar haplotypes, and regardless of UMI collision, low *ρ* may over-split (overcount in UMI; false positive in haplotype) PCR errors. Automated selection of *ρ* worked well for simulated and V1V2 data. It failed for V3, where error and true count distributions highly overlap because of high UMI collision and low amplification. In this case, manual selection based on the observed UMI frequencies worked. We also demonstrated that DAUMI has superior performance for a range of reasonable *ρ*. It may be possible to use cross-validation in large datasets or control variants to calibrate *ρ* or adaptively select distinct *ρ*_*sk*_ for each UMI/haplotype combination, perhaps as a simple function of ***u***_*s*_, ***h***_*k*_ abundance or the proximity and abundance of other observed molecules. Such attempts are increasingly complex approximations of an explicit PCR model.

DAUMI is also affected by the initial choice for the candidate UMI 𝒰 and haplotype ℋ sets. If candidates are known a priori, DAUMI works very well (Supplementary Material §S4), but most applications will have no prior knowledge of ℋ. To strike a good balance betwen sensitivity and specificity, we used AmpliCI [38] to remove the most likely error sequences. A side effect of our use of AmpliCI is that each true sample sequence must be observed with the same UMI at least twice to be considered. Singleton discard is a blunt way to increase agreement across subsets, as shown for Calib, UMI-tools and Starcode-umi, which do not discard singletons by default (Table 3 vs. Supplementary Table S6), The resulting UMI counts are biased downward, which can be corrected by estimating the PCR amplification process from the surviving replicates observed in the data [40], but the discarded haplotypes cannot be recovered. In low complexity samples with low amplification or high downsampling, multiple singleton or low-frequency UMIs may be linked to the same sampled molecule. DAUMI’s poorer performance on V1V2 when restricting to haplotypes with deduplicated abundance ≥ 2 (Table S6) is due to AmpliCI discard of barely amplified UMI-tagged haplotypes. AmpliCI uses more sophisticated logic to discard haplotypes than other tools, so its sensitivity can be increased if desired (see options `−−diagnostic` and `−−abundance`), but DAUMI defaults are set to work on varied datasets and without posthoc filtering.

As a flexible probabilistic model, it is possible to imagine extensions of DAUMI to additional use cases. Overpopulation of the candidate UMI set 𝒰 could be ameliorated by penalizing the relative abundances *η*_*s*_ of candidates ***u***_*s*_ ∈ 𝒰 in addition to the currently sparsified transition parameters in matrix ***Γ***. In fact, *η*_*s*_ is the product of the PCR amplification process, which could be explicitly modeled to detect early PCR errors or other known sources of variation in UMI abundance, such as biased nucleotide usage. The transition probabilities in ***Γ*** that link UMIs ***u***_*s*_ to haplotype ***h***_*k*_ could also be elaborated to model selection of a transcript followed by random selection of a fragmentation point within that transcript, as utilized in some scRNA-seq protocols [67]. Finally, the DAUMI framework is not limited to Illumina sequence data and could also, with a carefully chosen error model and a good initialization, be applied to technology that generates long reads.

