## Supplementary Material for "Accurate Estimation of Molecular Counts from Amplicon Sequence Data with Unique Molecular Identifiers"


June 12, 2022

### S1 EM Algorithm

We use an EM algorithm to estimate UMI proportions $\boldsymbol{\eta}$ and the transition parameters $\boldsymbol{\Gamma}$ that link UMIs to sample sequences. Conditional on the set of $N$ UMI candidates in $\mathcal{U}$ and $K$ haplotype candidates in $\mathcal{H}$, the complete data penalized log likelihood function for both the observed data $\mathcal{R}$ and $\mathcal{B}$ and unobserved hidden variables $\boldsymbol{Z}$ is

$$l(\boldsymbol{\eta}, \Gamma, \mid \mathcal{R}, \mathcal{B}, \boldsymbol{Z}) = \sum_{i=1}^{n}\sum_{s=1}^{N}\sum_{k=1}^{K}\left\{\log\left[\eta_s \Pr(\boldsymbol{b}_i \mid \boldsymbol{u}_s)\gamma_{sk}\Pr(\boldsymbol{r}_i \mid \boldsymbol{h}_k)\right]\right\}^{\mathbb{1}_{\{Z_{i1}=\boldsymbol{u}_s, Z_{i2}=\boldsymbol{h}_k\}}} - \rho\mathcal{J}(\Gamma). \tag{S1}$$

Define the conditional expectation of the hidden data $\boldsymbol{Z}_i = (Z_{i1}, Z_{i2})$ given the observed data $(\boldsymbol{b}_i, \boldsymbol{r}_i)$ and computed with the current parameter estimates $\boldsymbol{\eta}^{(t)}$ and $\Gamma^{(t)}$,

$$e_{isk}^{(t)} = \Pr(Z_{i1} = \boldsymbol{u}_s, Z_{i2} = \boldsymbol{h}_k \mid \boldsymbol{r}_i, \boldsymbol{b}_i; \boldsymbol{\eta}^{(t)}, \Gamma^{(t)}).$$

The EM algorithm iterates between an E step, where the expectations $e_{isk}^{(t)}$ are computed, and an M step, where the model parameters are estimated to maximize equation (S1), with $e_{isk}^{(t)}$ substituted for $\mathbb{1}_{\{Z_{i1}=\boldsymbol{u}_s, Z_{i2}=\boldsymbol{h}_k\}}$. Iterations will continue until convergence, as defined in the main text, to a local maximum of Eq. (1). To simplify the notation in the following, we drop the superscripts $(t)$ indicating the $t$th EM iteration, but each of $e_{isk}$, $\xi_{sk}$, $\lambda_s$, $\phi_{sk}$, some to be defined below, as well as the parameters $\boldsymbol{\eta}, \boldsymbol{\Gamma}$ and their updates $\hat{\boldsymbol{\eta}}, \hat{\boldsymbol{\Gamma}}$, will depend on the iteration $t$.

*E step:*

For the $i$th read, $e_{isk}$ is, by Bayes rule,

$$e_{isk} = \frac{\eta_s \Pr(\boldsymbol{b}_i \mid \boldsymbol{u}_s)\gamma_{sk} \Pr(\boldsymbol{r}_i \mid \boldsymbol{h}_k)}{\sum_{\boldsymbol{u}_s \in \mathcal{U}, \boldsymbol{h}_k \in \mathcal{H}} \eta_s \Pr(\boldsymbol{b}_i \mid \boldsymbol{u}_s)\gamma_{\boldsymbol{u}_s, \boldsymbol{h}_k} \Pr(\boldsymbol{r}_i \mid \boldsymbol{h}_k)},$$

where all probabilities are given in the main text and parameters are replaced with their current estimates.

*M step:*

Parameter $\boldsymbol{\eta}$ can be updated with

$$\hat{\eta}_s = \frac{1}{n} \sum_{i=1}^{n} \sum_{k=1}^{K} e_{isk}, \tag{S2}$$

while the $s$th row $\boldsymbol{\gamma}_s$ of transition matrix $\boldsymbol{\Gamma}$ can be updated by maximization of

$$\sum_{i=1}^{n} \sum_{k=1}^{K} e_{isk} \log(\gamma_{sk}) - \rho \sum_{k=1}^{K} \log(1 + \gamma_{sk}/\omega) - \lambda_s \left( \sum_{k=1}^{K} \gamma_{sk} - 1 \right), \tag{S3}$$

where $\lambda_s$ is the Lagrange multiplier to impose the sum constraint for transitions out of $\boldsymbol{u}_s$. In the absence of a penalty ($\rho = 0$), the MLE is

$$\hat{\gamma}_{sk} = \frac{\xi_{sk}}{\sum_{k=1}^{K} \xi_{sk}},$$

where $\xi_{sk} = \sum_{i=1}^{n} e_{isk}$ is the expected number of molecules where UMI $\boldsymbol{u}_s$ is attached to molecule $\boldsymbol{h}_k$. For positive $\rho$, the maximum penalized-likelihood estimators (MPLEs) of $\boldsymbol{\gamma}_s$ are $\hat{\gamma}_{sk} = 0$ when $\xi_{sk} = 0$ and solutions of the score functions,

$$\frac{\xi_{sk}}{\gamma_{sk}} - \frac{\rho}{\omega + \gamma_{sk}} - \lambda_s = 0,$$

for each $k$ with $\xi_{sk} > 0$. Yin *et al.* [1] prove the solution to these score functions exists and maximizes the objective function when (a) at least one $\xi_{sk} > \rho$ and (b) $|\xi_{sk} - \rho| > \omega$ for all $k$. Under these conditions, $\hat{\gamma}_{sk}$ is one of the roots

$$\frac{\phi_{sk} \pm \sqrt{\phi_{sk}^2 + 4\omega\lambda_s\xi_{sk}}}{2\lambda_s},$$

where $\phi_{sk} = \xi_{sk} - \omega\lambda_s - \rho$ and $\lambda_s$ is the root of implicit equation

$$\sum_{k=1}^{K} \hat{\gamma}_{sk} = 1. \tag{S4}$$

In practice, we set $\omega$ to a very small value (default: $10^{-20}$) for LASSO-like regularization and apply the Newton-Raphson algorithm to find the root $\lambda_s$ initialized as $\max_{0 \leq k \leq K} \xi_{sk} - \rho$.

Since $\omega$ is very small, it is rare that condition (b) is not met, especially when $\rho$ is chosen not to be an integer. Condition (a) is not met (all $\xi_{sk} < \rho$) when (c) $\boldsymbol{u}_s$ is an error UMI incorrectly included in $\mathcal{U}$ or (d) $\boldsymbol{u}_s$ is a valid UMI that was only weakly amplified, probably with early-cycle PCR error(s) to haplotype(s) $\boldsymbol{h}_k$ identical or similar to haplotypes included in $\mathcal{H}$. The first problem (c) may be resolved by applying a penalty to $\eta_s$, though we did not attempt such a solution in DAUMI since we took other precautions to limit the inclusion of false UMIs in $\mathcal{U}$. The second problem (d) is difficult to resolve without properly modeling the PCR error and amplification process. DAUMI and all compared methods do not model PCR and cannot distinguish early-cycle PCR errors from legitimate variation in the sample. Instead, these methods rely on the fact that early-cycle PCR errors are exceptionally rare compared to late-cycle PCR errors, because there is far more opportunity for error after the template has been geometrically amplified. Thus, these quantification methods detect and discard most PCR and sequencing errors, but inevitably leave a few PCR errors. In these few remaining cases, we set $\hat{\gamma}_{sk} = \mathbb{1}_{\{k=\hat{k}\}}$ for $\hat{k} = \arg\max_k \xi_{sk}$. As long as the overall M step improves equation (S1) at each iteration, the EM algorithm is still guaranteed to converge [2]. We check for this condition and never encountered a problem in any of the real or simulation datasets analyzed.

### S2 Automated Selection of Penalty Parameter $\rho$

To automate selection of penalty parameter $\rho$, we propose a simple model for the observed abundance distribution $p_{\text{UMI}}(x)$ of UMIs in the dataset. This distribution is a mixture of true UMIs and error UMIs generated by PCR and sequencing errors. Our goal is to identify a threshold that distinguishes true and error UMIs.

We assume molecular abundance follows a stochastic Galton-Watson branching process [3], which is often used to model the PCR amplification process [4]. The model assumes each molecule is independently duplicated with amplification efficiency $\chi \in (0, 1)$ per PCR cycle. Let $\epsilon$ denote the PCR error rate, a small probability that a molecule is copied with at least one error. The number of *error-free* copies $X_t$

of a true UMI without collision after the $t$th PCR cycle is given by the stochastic recursion

$$X_0 = 1, \quad X_t = X_{t-1} + \mathrm{Bin}(X_{t-1}, [1-\epsilon]\chi),$$

where $[1-\epsilon]\chi$ is the *error-free* amplification efficiency. After $\sigma$ cycles of PCR, the molecules are sampled and sequenced to produce observed count

$$X_{\sigma+1} \sim \mathrm{Bin}(X_\sigma, [1-\delta]\iota),$$

where $\iota$ is the sampling rate and $\delta$ is the sequencing error probability.

The probability $p_{\mathrm{BP}}(x; \sigma, \chi, \epsilon, \iota, \delta)$ of $x$ faithful copies of a molecule after $\sigma$ cycles of PCR amplification followed by sequencing is not analytically available, but can be computed by numerical methods [5]. Assuming all PCR amplification errors are unique, the abundance of an error UMI depends only on the random PCR cycle $T$ in which it originated and not also on the amplification of similar molecules in a neighborhood around it. Then, using the law of total probability, the probability of observing $x$ copies of an error UMI is

$$\sum_{1 \le t \le \sigma} p_{\mathrm{BP}}(x; \sigma - t, \chi, \epsilon, \iota, \delta) \Pr(T = t) + \mathbb{1}_{\{x=1\}} \Pr(T = \sigma + 1),$$

where $\Pr(T = t)$ is the probability of a PCR error in the $t$th cycle or a sequencing error when $T = \sigma + 1$.

Overall, the abundance of a UMI follows a mixture distribution

$$p_{\mathrm{UMI}}(x) = p_{\mathrm{BP}}(x; \sigma, \chi, \epsilon, \iota, \delta) \Pr(T = 0) + \sum_{1 \le t \le \sigma} p_{\mathrm{BP}}(x; \sigma - t, \chi, \epsilon, \iota, \delta) \Pr(T = t) + \mathbb{1}_{\{x=1\}} \Pr(T = \sigma + 1),$$

where we have included error-free amplification of original molecules present at $T = 0$. In practice, we take set $\iota = 1$ when calculating $p_{\mathrm{UMI}}(x)$. In our experience, $\iota$ is substantially confounded with $\chi$ and $\sigma$, when the latter is unknown. Setting $\iota = 1$ constrains the parameters sufficiently to produce good estimates, as assessed on simulation data. Assuming UMIs are unique prior to amplification (no collision), we have

$$\Pr(T = t) \propto \begin{cases} 1 & t = 0 \\ \epsilon(1+\chi)^{t-1} & 1 \le t \le \sigma \\ \delta(1+\chi)^{\sigma} & t = \sigma + 1, \end{cases}$$

so $p_{\mathrm{UMI}}(x)$ is a function of the parameters $\{\chi, \sigma, \epsilon, \delta\}$, which we can optimize to match the observed UMI abundance distribution.

There is no analytical solution to estimate the parameters $\{\chi, \sigma, \epsilon, \delta\}$, and the presence of unmodeled artefacts, like collision, in the right tail of the abundance distribution makes numerical maximum likelihood estimation difficult. Instead, we right-truncate the data at count $\tau$ and select parameters that minimize the Kolmogorov-Smirnov (KS) statistic [6] for conditional distribution

$$p_{\text{UMI}}(x \mid x \leq \tau) = \Pr(X_{\omega+1} = x \mid X_{\omega+1} \leq \tau).$$

From the left, the first mode of the observed UMI abundance distribution represents errors, while the second and possibly additional modes represent PCR amplified products [3, 4]. To estimate the parameters of the PCR process, it is important to retain information about PCR amplification by truncating the distribution to the right of the second mode. We use R package `multimode` [7] to find the mode of each distribution after removing sequencing errors with AmpliCI (Figures S3 and S5). Given the conditional cumulative distribution function $F(x) = \sum_{y=0}^{x} p_{\text{UMI}}(y \mid y \leq \tau)$ and the empirical distribution function $F_{n(\tau)}(x) = \frac{1}{n(\tau)} \sum_{s=1}^{n(\tau)} \mathbb{1}_{\{X_s \in [0,x]\}}$ for $n(\tau)$ observed UMIs $b_s$ with abundance $X_s = \sum_{i=1}^{n} \mathbb{1}_{\{\boldsymbol{b}_i = \boldsymbol{b}_s\}} \leq \tau$, the KS statistic is the largest absolute difference between these two distributions, $\sup_{x \in \{0,1,2,\ldots\}} |F_{n(\tau)}(x) - F(x)|$. Specifically, we use a grid search over $\tau \in \{100, 150, 200, 300\}$, $\chi \in \{0.05, 0.10, \ldots, 1\}$, $\sigma \in \{6, 7, \ldots, 21\}$, $\epsilon \in \{0.0005, 0.0010, \ldots, 0.005\}$ and $\delta \in \{0.005, 0.006, \ldots, 0.014\}$. The ranges are selected based on the values observed in real experiments [8, 3, 9, 10]. Rather than an exhaustive search over the entire grid, we implement a block relaxation algorithm [11], alternating between optimization of $(\chi, \sigma)$ while holding $(\epsilon, \delta)$ fixed and vice versa until the KS statistic converges with the relative change between iterations $< 10^{-4}$ for each possible value of $\tau$. The estimated $\hat{\sigma}$ is not necessarily the actual number of experimental PCR cycles, but an *effective* number of PCR amplification cycles under our "ideal" model of PCR amplification.

The remaining challenge is to choose a threshold $\rho$ that will remove most error UMIs without discarding true UMIs. In practice, we choose $\rho$ to be the 5th percentile of the fitted observed abundance distribution $p_{\text{BP}}(x; \hat{\sigma}, \hat{\chi}, \hat{\epsilon}, 1, \hat{\delta})$ for true UMIs, which implies an estimated five percent of true variants will have observed abundance below the threshold $\rho$. Table S1 shows the fitted model and selected $\rho$ for each $\tau \in \{100, 150, 200, 300\}$ in all test datasets. For most datasets, our strategy works well and helps to select appropriate $\rho$ (Figure S2 and S4). On V1V2 dataset, the strategy also selects the optimum $\rho$ (Table S5), but the strategy failed on the V3 dataset, always choosing $\rho = 1$, no matter the truncation point $\tau$. There is no obvious second mode in these data, either because there was very inefficient amplification, we estimate $\hat{\chi} = 0.05$, or severe downsampling at sequencing. Both scenarios limit UMI replication and inhibit all UMI-based quantification methods. At the same time, these data display some very highly replicated UMIs, perhaps because of high rates of UMI collision or highly stochastic amplification rates [9]. For all these reasons, it is difficult to pick a threshold to separate errors from real variants. For such recalcitrant data, we suggest visual assessment of the observed UMI abundances and caution in the interpretation of results. Inspired by the values of $\rho$ selected automatically for other datasets and comparing observed UMI abundance values (Figure S4) and observed UMI abundance after error removal by AmpliCI (Figure S5), we manually choose $\rho = 20$ for the V3 dataset. For V1V2, $\rho$ was selected independently for each subset by the automatic procedure. For V3, $\rho = 10$ was selected for both subsets, half the value chosen by visual assessment for the whole dataset. Selection of $\rho$ generally has little impact on DAUMI performance unless $\rho$ is very small (Supplementary Table S5).

### S3 Tuning Run Parameters on Simulated Datasets

As just demonstrated for selection of $\rho$, it is important and sometimes difficult to set appropriate values for run parameters; the default values are not always optimal. To verify the optimality of the default parameters used in the quantification methods studied in this work, we ran Calib, UMI-tools and DAUMI under different parameter settings. There are no run parameters for the Naïve method, and tuning Starcode-umi was deemed too difficult as there are at least five parameters and no guidance to set them.

Table S2 shows the results on simulation data while varying run parameters. Specifically, we try edit distance $d = 1$ or $d = 2$ for UMI-tools, vary $\rho$ for DAUMI, and try six parameter settings for Calib as suggested by the authors [12]. UMI-tools performs better with the default edit distance $d = 1$, though the number of false positives slightly increases. While Calib's default parameter values did not always produce the best results, the effect of different parameter settings was minimal. We show that DAUMI at default settings always outperformed other algorithms at their optimal parameter settings, even though default $\rho$ does not always achieve the best performance.

### S4 Abundance Estimation for Known Variants

In some cases, the set of possible haplotypes is known a priori [13]. To mimic this situation, we set $\mathcal{H}$ to the true 25 haplotypes for simulations 1–4. This experiment also allows us to explore DAUMI abundance estimation with less chance for false positives, since only a few of the 25 true haplotypes are not sampled in the data and could be incorrectly included as false positives. Table S3 shows that performance does increase as compared to the performance in Table S2, where the set $\mathcal{H}$ is not known. DAUMI can accurately estimate deduplicated abundance on datasets *without* UMI collision (Simulations 1–2) as expected, and there is little difference as $\rho$ varies. However, for datasets *with* UMI collision (Simulations

3–4), DAUMI underestimates haplotype deduplicated abundance (regression slope $b < 1$), especially as $\rho$ increases, because then even legitimate UMI to haplotype linkages with transition probability $\gamma_{sk} > 0$ are eliminated by the penalty. For Simulation 4 with low PCR efficiency, the underestimation is particularly sensitive to $\rho$, presumably because there are more true molecules failing to amplify well. When two haplotypes are assigned to the same UMI and the number of assigned reads to one haplotype is below the threshold $\rho$, DAUMI will eliminate the less observed haplotype as a likely error, leading to an underestimation of haplotype abundance. However, it is not a good idea to simply set $\rho$ very low, since then DAUMI will retain false linkages between UMIs and haplotypes. There may be weak evidence for UMI/haplotype combinations $(\boldsymbol{u}_s, \boldsymbol{h}_k)$ that do not actually exist in the sample if there is at least one observed read $(\boldsymbol{b}_i, \boldsymbol{r}_i)$ plausibly generated from $(\boldsymbol{u}_s, \boldsymbol{h}_k)$. For example, DAUMI identifies two false positives for Simulation 3 and one false positive for Simulation 4 with $\rho = 0.01$. These are haplotypes included in $\mathcal{H}$ that were not actually sampled. When the haplotype set $\mathcal{H}$ is not constrained by a known truth, there will be even more weak linkages that survive when $\rho$ is set too low.

### S5 Extracting UMIs from V1V2 Dataset

The reverse reads of the V1V2 dataset contain a 52–55nt technical sequence at the 5' end, starting with 0–3 random nucleotides, an 18nt adapter, a 9nt UMI, and a 25nt primer with two ambiguous nucleotides, one Y and another R, used to amplify the target region in the HIV *env* gene. Our goal is to extract the 9bp UMI, discard the rest of the technical sequence, and recover the approximately 250nt sampled sequence.

We assume there are no indel errors in the technical sequence of each read, so the adapter sequence starts at read position 1, 2, 3, 4, or is not present at all. Let $Z_{i1} \in \{0, 1, 2, 3, 4\}$ be the unknown number of random nucleotides at the start of read $\boldsymbol{X}_i$ or $Z_{i1} = 4$ when the technical sequence is not present. Further, let $Z_{i2} \in \{0, 1, 2, 3\}$, defined only when $Z_{i1} < 4$, be the unknown state of the *unambiguous* primer sans UMI with the Y and R nucleotides resolved. We assume $Z_{i1}$ and $Z_{i2}$ are independent and define $\eta_k = \Pr(Z_{i1} = k)$ and $\zeta_l = \Pr(Z_{i2} = l \mid Z_{i1} < 4)$. The bivariate $\boldsymbol{Z}_i = (Z_{i1}, Z_{i2})$ is unobserved.

For the observed data, dropping read index $i$, let

$$p_{jx}(\boldsymbol{z}) = \Pr(X_j = x \mid \boldsymbol{Z} = \boldsymbol{z})$$

be the probability of read nucleotide $x$ at position $j$ given $\boldsymbol{Z} = \boldsymbol{z}$. Read position $j$ can index the 0–3 nucleotides in the 5' random leader, nucleotides in the UMI, either of two ambiguous positions in the primer, other unambiguous nucleotides in the technical sequence or the sample sequence. Assume the random 5' leader, the UMI and the sampled sequence are adequately modeled as independently and identically distributed nucleotides within class, but allow the nucleotide composition to vary: $\boldsymbol{q}_r = (q_{r\mathrm{A}}, q_{r\mathrm{C}}, q_{r\mathrm{G}}, q_{r\mathrm{T}})$ for the random nucleotides in the 5' leader and UMI and $\boldsymbol{q}_s = (q_{s\mathrm{A}}, q_{s\mathrm{C}}, q_{s\mathrm{G}}, q_{s\mathrm{T}})$ for the sample sequence. At all other sites, assume errors are independent, but not equally likely across sites. Let $\delta_j$ be the probability of an error-free nucleotide at read position $j$. Given an error, let $\gamma_{\mathrm{N}_1\mathrm{N}_2}$ be the probability that true nucleotide $\mathrm{N}_1$ is misread as read nucleotide $\mathrm{N}_2$. Define the index sets $\mathcal{V} = \{19, 20, \ldots, 27\}$ for the UMI indices, $\mathcal{T} = \{1, 2, \ldots, 52\}\backslash\{\mathcal{V}, 35, 42\}$ for the unambiguous technical sequence, and $\mathcal{S} = \{53, 54, \ldots\}$ for the sample sequence. Let $\boldsymbol{D} = (D_1, D_2, \ldots, D_{52})$ be the 52 nucleotides in the technical sequence, excluding the 5' leader, Nucleotides $D_{35} = \mathrm{R}$ and $D_{42} = \mathrm{Y}$ are ambiguous, so let $R_l, Y_l \in \{\mathrm{A}, \mathrm{C}, \mathrm{G}, \mathrm{T}\}$ be the resolved nucleotides when $Z_2 = l$, and $\mathbb{1}_C(x) = \mathbb{1}_{\{x \in C\}}$ indicate the event $x \in C$. Still dropping read index $i$, we have

$$
\begin{aligned}
p_{jx}(\boldsymbol{z}) &= \Pr(X_j = x | \boldsymbol{Z} = \boldsymbol{z}) \\
&= (q_{sx})^{\mathbb{1}_{\{z_1=4\}}} \Bigg[ (q_{bx})^{\mathbb{1}_{\{j \le z_1\}} + \mathbb{1}_{\mathcal{V}}(j-z_1)} (q_{sx})^{\mathbb{1}_{\mathcal{S}}(j-z_1)} \\
&\quad \left( \delta_j^{\mathbb{1}_{\{x=Y_l\}}} [(1-\delta_j)\gamma_{Y_l x}]^{\mathbb{1}_{\{x \ne Y_l\}}} \right)^{\mathbb{1}_{\{35\}}(j-z_1)} \left( \delta_j^{\mathbb{1}_{\{x=R_l\}}} [(1-\delta_j)\gamma_{R_l x}]^{\mathbb{1}_{\{x \ne R_l\}}} \right)^{\mathbb{1}_{\{42\}}(j-z_1)} \\
&\quad \left( \delta_j^{\mathbb{1}_{\{x=D_{(j-z_1)}\}}} [(1-\delta_j)\gamma_{D_{(j-z_1)}x}]^{\mathbb{1}_{\{x \ne D_{(j-z_1)}\}}} \right)^{\mathbb{1}_{\mathcal{T}}(j-z_1)} \Bigg]^{\mathbb{1}_{\{z_1 \le 3\}}} .
\end{aligned}
$$

Let $\boldsymbol{\delta} = (\delta_1, \delta_2, ...)^T$, $\boldsymbol{\gamma} = (\gamma_{\mathrm{AC}}, \gamma_{\mathrm{AG}}, \gamma_{\mathrm{AT}}, \gamma_{\mathrm{CA}}, \gamma_{\mathrm{CG}}, \gamma_{\mathrm{CT}}, \gamma_{\mathrm{GA}}, \gamma_{\mathrm{GC}}, \gamma_{\mathrm{GT}}, \gamma_{\mathrm{TA}}, \gamma_{\mathrm{TC}}, \gamma_{\mathrm{TG}})$, $\boldsymbol{\eta} = (\eta_0, \eta_1, \ldots, \eta_4)$, and $\boldsymbol{\zeta} = (\zeta_0, \zeta_1, \zeta_2, \zeta_e)$. Then our unknown parameter vector is $\boldsymbol{\theta} = (\boldsymbol{\delta}^T, \boldsymbol{\gamma}^T, \boldsymbol{q}_u^T, \boldsymbol{q}_s^T, \boldsymbol{\eta}^T, \boldsymbol{\zeta}^T)^T$. Finally, if there are $n$ reads, the length of read $i$ is $J_i$, all reads are independent and all nucleotides within reads are conditionally independent, then the complete data likelihood is

$$
L_C(\boldsymbol{\theta} \mid \boldsymbol{X}, \boldsymbol{Z}) = \prod_{i=1}^{n} \Pr(\boldsymbol{X}_i = \boldsymbol{x}_i, \boldsymbol{Z}_i = \boldsymbol{z}_i) = \prod_{i=1}^{n} \prod_{k=0}^{4} \prod_{l=0}^{3} \Big[ \Pr(Z_{i1} = k, Z_{i2} = l) \prod_{j=1}^{J_i} p_{X_{ij}}(k, l) \Big]^{\mathbb{1}_{\{Z_{i1}=k, Z_{i2}=l\}}} .
$$

In the E step, we need to compute $\mathbb{E}[\ln L_C(\boldsymbol{\theta} \mid \boldsymbol{X}, \boldsymbol{Z}) \mid \boldsymbol{X}]$, but since the complete data log likelihood is linear and the reads are independent, it amounts to computing

$$
\begin{aligned}
e_{ikl} &:= \Pr(Z_{i1} = k, Z_{i2} = l \mid \boldsymbol{X}_i) \\
&\propto \eta_k \zeta_l^{\mathbb{1}_{\{k<4\}}} \prod_{j=1}^{J_i} p_{jX_{ij}}(l, k),
\end{aligned}
$$

for each $i \in \{1, 2, \ldots, n\}$, $k \in \{0, 1, 2, 3\}$, and $l \in \{0, 1, 2, 3\}$. When $k = 4$, $l$ is undefined, and we have

$e_{i4} = \Pr(Z_{i1} = 4 \mid \boldsymbol{X}_i)$. The update equations in the M step are

$$\eta_k = \frac{n_{\eta k}}{\sum_l n_{\eta l}} \qquad \zeta_l = \frac{n_{\zeta l}}{\sum_{k=0}^{3} n_{\zeta k}}$$
$$q_{ux} = \frac{n_{ux}}{\sum_y n_{uy}} \qquad q_{sx} = \frac{n_{sx}}{\sum_y n_{sy}}$$
$$\delta_j = \frac{n_{j0}}{n_{j0} + n_{j1}} \qquad \gamma_{yx} = \frac{n_{yx}}{\sum_z n_{yz}},$$

with expected counts

$$n_{\eta k} = \begin{cases} \sum_{i=1}^{n} e_{ikl} & k < 4 \\ \sum_{i=1}^{n} e_{ik} & k = 4 \end{cases}$$
$$n_{\zeta l} = \sum_{i=1}^{n} e_{ikl}, k < 4$$
$$n_{ux} = \sum_{i=1}^{n}\sum_{k=0}^{3}\sum_{l=0}^{3} e_{ikl} \sum_{j=1}^{J_i} \mathbb{1}_{\{X_{ij}=x\}}(\mathbb{1}_{\{j\le k\}} + \mathbb{1}_{\mathcal{V}}(j-k))$$
$$n_{sx} = \sum_{i=1}^{n}\Big[\sum_{k=0}^{3}\sum_{l=0}^{3} e_{ikl} \sum_{j=k+52}^{J_i} \mathbb{1}_{\{X_{ij}=x\}} + e_{i4}\sum_{j=1}^{J_i} \mathbb{1}_{\{X_{ij}=x\}}\Big]$$
$$n_{j0} = \sum_{i=1}^{n}\sum_{k=0}^{3}\sum_{l=0}^{3} e_{ikl}\mathbb{1}_{\{X_{i,j+k}=D_j\}}\left[\mathbb{1}_{\mathcal{T}}(j-k) + \mathbb{1}_{\{35\}}(j-k) + \mathbb{1}_{\{42\}}(j-k)\right]$$
$$n_{j1} = \sum_{i=1}^{n}\sum_{k=0}^{3}\sum_{l=0}^{3} e_{ikl}\mathbb{1}_{\{X_{i,j+k}\neq D_j\}}\left[\mathbb{1}_{\mathcal{T}}(j-k) + \mathbb{1}_{\{35\}}(j-k) + \mathbb{1}_{\{42\}}(j-k)\right]$$
$$n_{yx} = \sum_{i=1}^{n}\sum_{k=0}^{3}\sum_{l=0}^{3} e_{ikl}\sum_{j=1}^{J_i}\mathbb{1}_{\{X_{i,j+k}=x,D_j=y\}}\left[\mathbb{1}_{\mathcal{T}}(j-k) + \mathbb{1}_{\{35\}}(j-k) + \mathbb{1}_{\{42\}}(j-k)\right].$$

We fit all 61,881 reads to this model, converting ambiguous N nucleotides (affecting just 170 reads) to A. We iterated the EM algorithm until the relative change in log likelihood was below 0.001. We dropped 18,861 reads shorter than the 52nt primer and 700 reads unlikely to contain the technical sequence, *i.e.* with $\Pr(Z_{i1} = 4 \mid \boldsymbol{X}_i) \geq 0.5$. We dropped three reads with posterior probability $\Pr(Z_{i1} = 4 \mid \boldsymbol{X}_i) < 0.5$, but $\arg\max_{k\in\{0,1,2,3\},l\in\{0,1,2,3\}} \Pr(Z_{i1} = k, Z_{i2} = l \mid \boldsymbol{X}_i) < \Pr(Z_{i1} = 4 \mid \boldsymbol{X}_i)$; these could have been retained but were lost to a small bug in our code. Since we noticed several reads had a poor match to the 25nt primer, we dropped another 8,763 reads where the log likelihood of the primer region only was smaller than $-100$ as likely technical artefacts. Finally, we dropped 25 surviving reads with ambiguous N nucleotides. For the 33,529 remaining reads, we assumed the 9nts starting at position $\hat{z}_{i1} + 18$ of the $i$th read constituted the UMI and the sampled molecule extended from $\hat{z}_{i1} + 52$ to the end of the read. Here, $\hat{z}_{i1} = \arg\max_{k\in\{0,1,2,3\}} \Pr(Z_{i1} = k \mid \boldsymbol{X}_i)$ is the most likely number of 5' leader nucleotides. The posterior probabilities used for screening were obtained from the $e_{ikl}, k < 4$ and $e_{i4}$ values computed in the last E step.

### S6 Supplementary Figures and Tables

Here we include all supplementary figures and tables that have been cited in the main text or in supplementary sections S1–S4. Figure S1 compares the main features of DAUMI and other methods. Figure S2, S3, S4, and S5 illustrate the observed UMI abundance distributions and the selected parameter $\rho$ in all simulated and real datasets. Table S1 provides detailed information about the achieved fits during selection of $\rho$. Table S2 and S3 show the performance of different methods with varying run parameters on the simulation datasets. Figure S6, S7, S8, and Table S4, S5, S6 show additional analysis results on the two real HIV datasets. Figure S7 provides additional evidence of UMI collision in the V3 dataset.

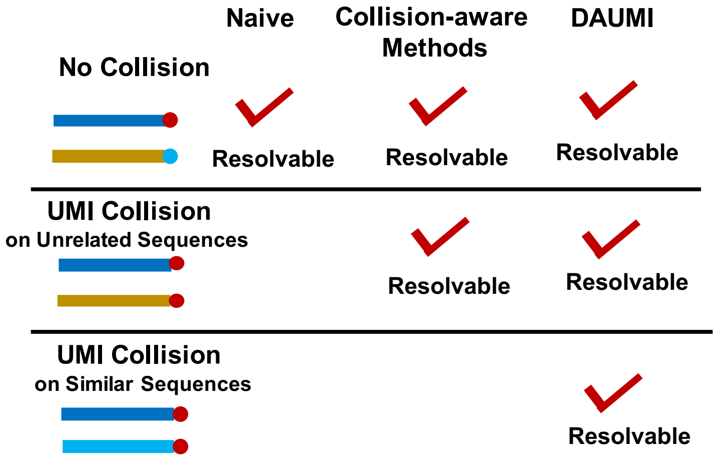


Figure S1: DAUMI can correctly resolve the origin of haplotypes even if there are UMI collisions.

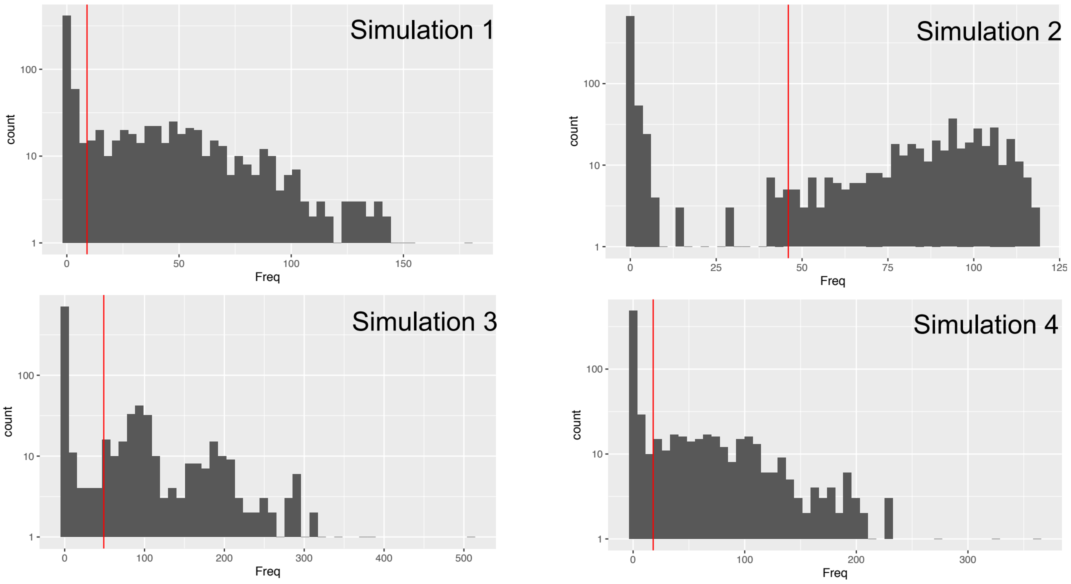


Figure S2: UMI observed abundance distribution of simulation datasets (errors uncorrected). Simulations 1 and 4 were simulated with PCR efficiency 0.5 in 10 cycles. Simulations 2 and 3 were simulated with PCR efficiency 0.9 in 7 cycles. There are UMI collisions in Simulations 3 and 4. The red vertical line is the $\rho$ selected by our proposed method.

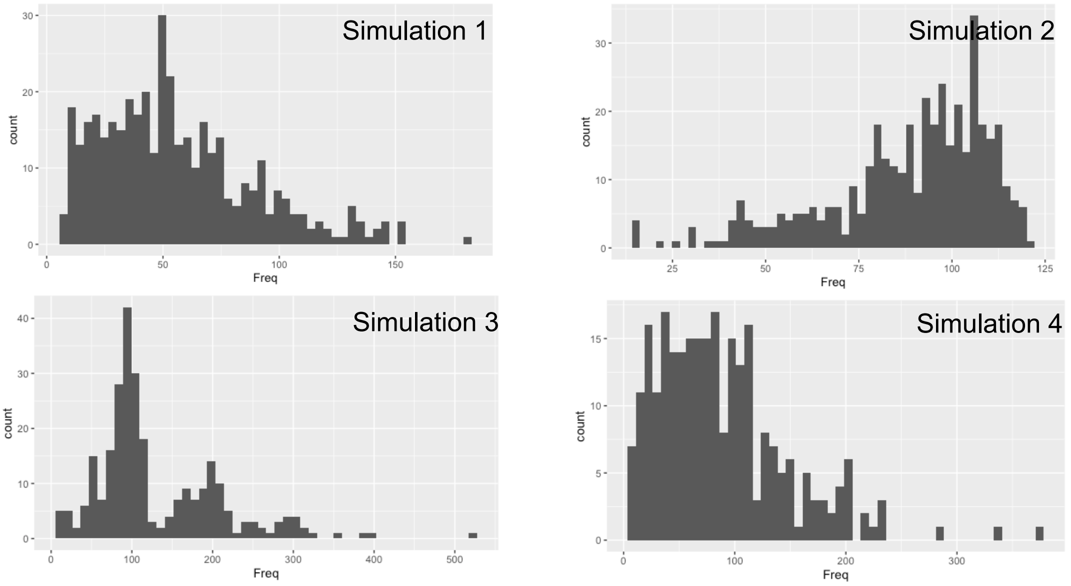


Figure S3: UMI abundance distribution of simulation datasets after removing sequencing errors with AmpliCI. See legend of Figure S2 for more details about the simulations.

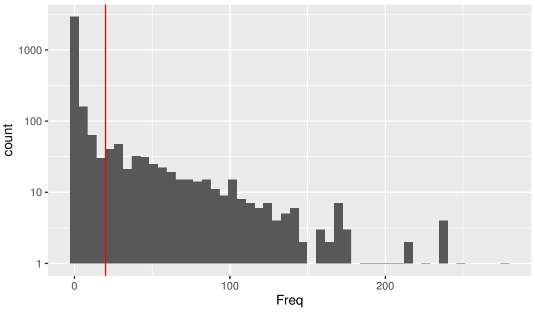


(a) VIV2

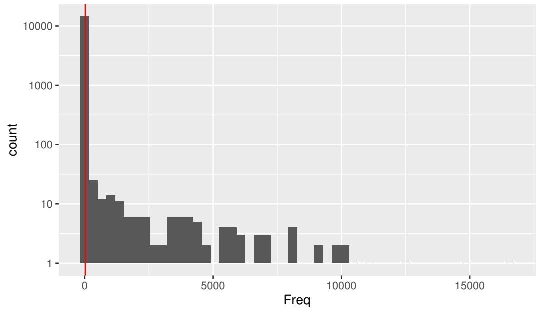


(b) V3

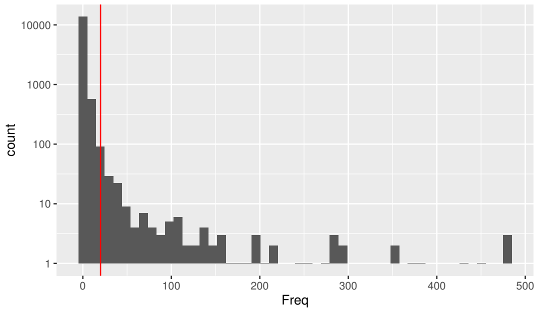


(c) V3 (Freq $\leq$ 500)

Figure S4: UMI observed abundance distribution of V1V2 and V3 datasets (errors uncorrected). V3 is highly right-skewed, so (c) focuses on the left tail for UMI with observed abundance at or below 500. The red vertical line is the $\rho$ selected either by our proposed method (V1V2) or by eye (V3).

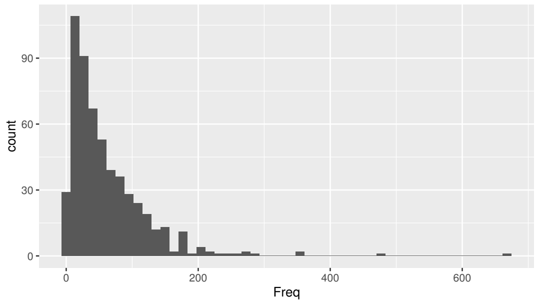


(a) VIV2

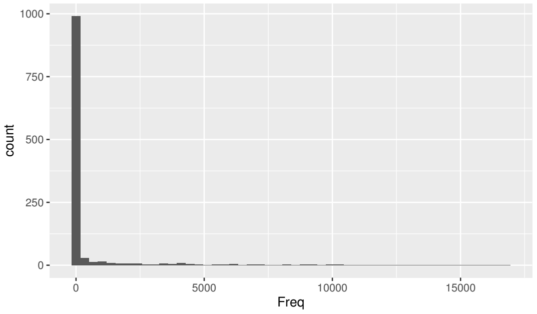


(b) V3

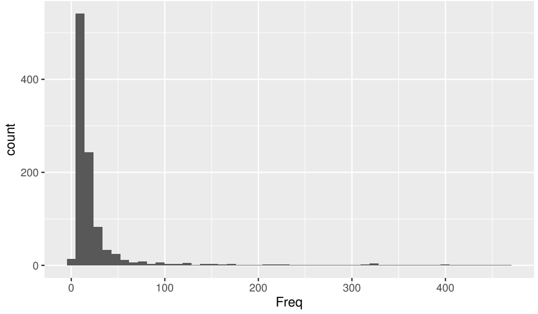


(c) V3 (Freq $\leq$ 500)

Figure S5: UMI abundance distribution of V1V2 and V3 after removing sequencing errors with AmpliCI. Again, (c) focuses on the left tail of the V3 distribution.

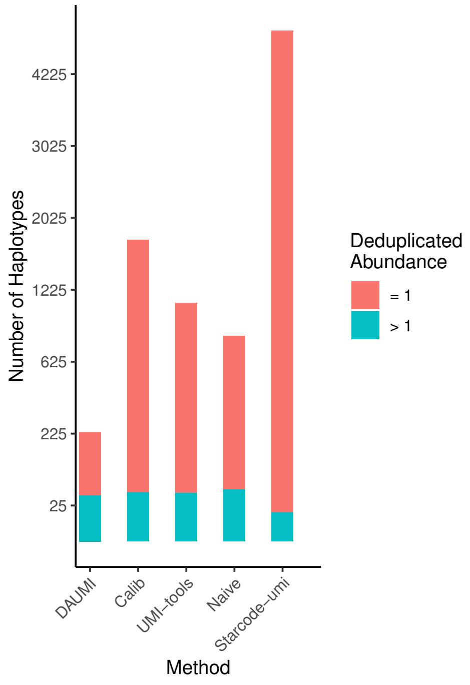


(a)

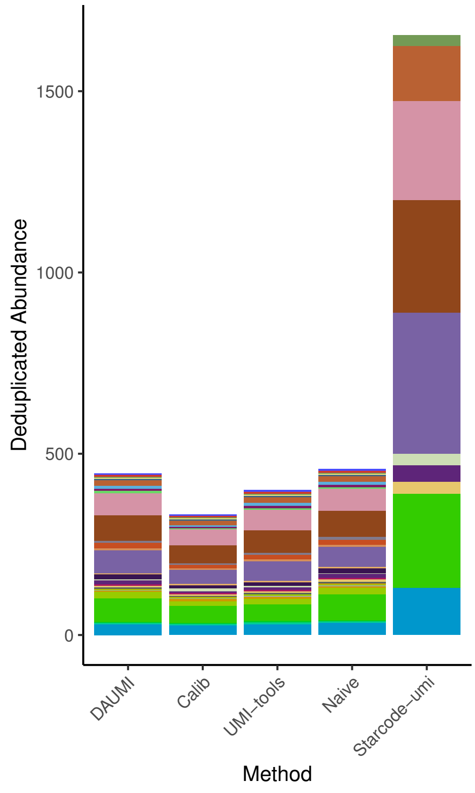


(b)

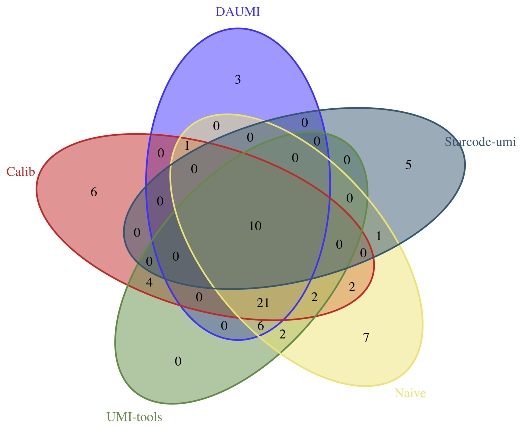


(c)

Figure S6: Results for V1V2 data [14] by DAUMI ($\rho = 20$), Calib, UMI-tools, Starcode-umi and Naïve methods. (a) Total number of recovered haplotypes, plotted on square root scale. (b) Estimated deduplicated abundance of the 31 haplotypes identified by all methods except Starcode-umi. (c) Venn Diagram of recovered haplotypes with deduplicated abundance $\geq 2$, made by VennDiagram R package (v1.6.20) [15].

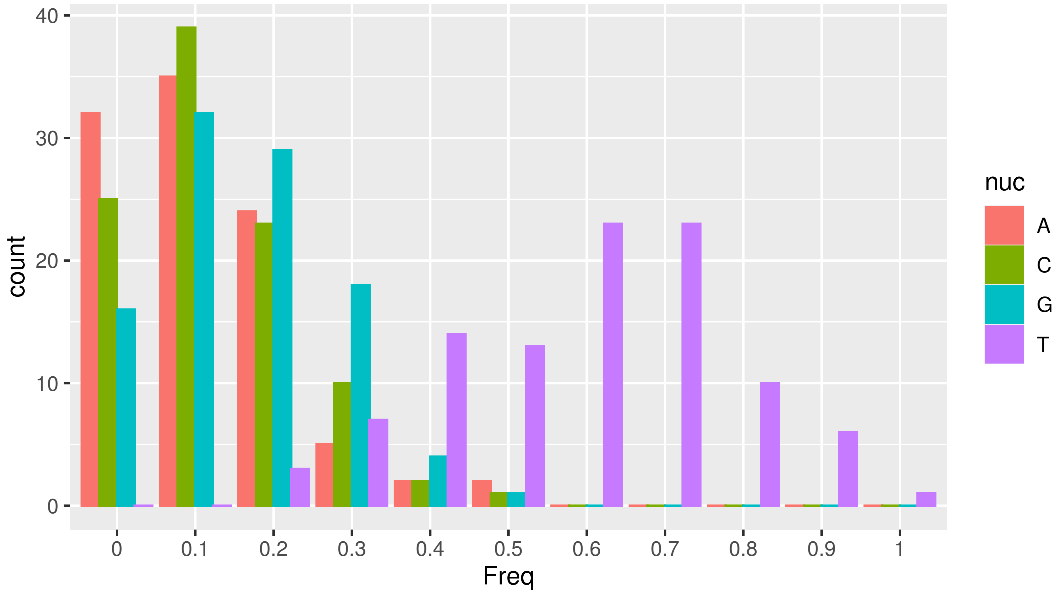


Figure S7: Nucleotide usage frequency per UMI in abundant UMIs of V3 dataset. The UMIs are of length 10nt in this dataset. We only take into account the top 100 UMIs, thus the total counts for each color should be equal to 100.

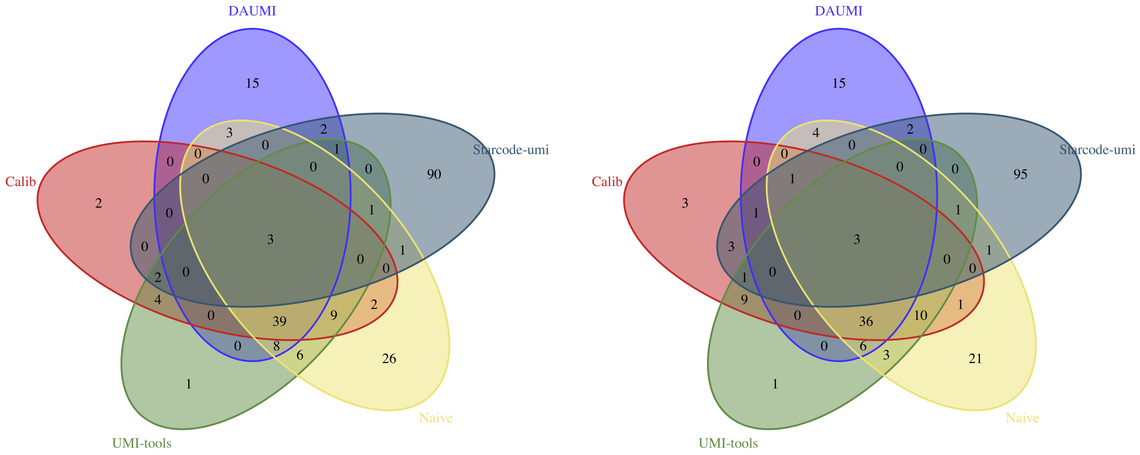


(a) Subset 1 (b) Subset 2

Figure S8: Venn diagrams comparing recovered haplotypes (with deduplicated abundance no less than two) on two subsets of V3 [16] by DAUMI ($\rho = 10$), Calib, UMI-tools, Starcode-umi and Naïve methods, made by VennDiagram R package (v1.6.20) [15]. Results of the first out of five random partitions is shown.

Table S1: Selection of right-tail truncation point $\tau$ when modeling UMI abundance distribution. To remove the unmodeled UMI collisions, we only retain UMI with observed counts $X \leq \tau$. Trunc.: truncation point $\tau$; Eff.: estimated PCR amplification efficiency $\hat{\chi}$; Cycles: estimated effective number $\hat{\sigma}$ of PCR cycles; PCR Err.: estimated probability $\hat{\epsilon}$ of PCR error; Seq. Err.: estimated probability $\hat{\delta}$ of sequencing error. Thres.: selected penalty parameter $\rho$. KS: Kolmogorov-Smirnoff goodness-of-fit test statistic, minimized to select the truncation point. Mode is the estimated mode of the observed abundance distribution after removing error UMIs with AmpliCI.

| **Datasets** | **Trunc.** $\tau$ | **Eff.** $\hat{\chi}$ | **Cycles** $\hat{\sigma}$ | **PCR Err.** $\hat{\epsilon}$ | **Seq. Err.** $\hat{\delta}$ | **Thres.** $\rho$ | **KS** |
|---|---|---|---|---|---|---|---|
| Simulation 1 | **100** | **0.45** | **11** | **0.0035** | **0.013** | **9** | **0.0194** |
| (mode: 45) | 150 | 0.45 | 11 | 0.0040 | 0.013 | 10 | 0.0213 |
| | 200 | 0.45 | 11 | 0.0040 | 0.013 | 10 | 0.0246 |
| | 300 | 0.45 | 11 | 0.0040 | 0.013 | 10 | 0.0249 |
| Simulation 2 | 100 | 0.75 | 8 | 0.0050 | 0.014 | 25 | 0.0511 |
| (mode: 102) | 150 | 0.90 | 7 | 0.0050 | 0.014 | 45 | 0.0307 |
| | 200 | 0.90 | 7 | 0.0050 | 0.014 | 45 | 0.0307 |
| | **300** | **0.90** | **7** | **0.0050** | **0.014** | **45** | **0.0307** |
| Simulation 3 | **100** | **0.95** | **7** | **0.0050** | **0.012** | **49** | **0.0145** |
| (mode: 96) | 150 | 0.65 | 10 | 0.0050 | 0.012 | 30 | 0.0443 |
| | 200 | 0.60 | 11 | 0.0035 | 0.010 | 34 | 0.0486 |
| | 300 | 0.50 | 12 | 0.0040 | 0.014 | 24 | 0.0361 |
| Simulation 4 | **100** | **0.60** | **10** | **0.0045** | **0.009** | **18** | **0.0201** |
| (mode: 66) | 150 | 0.60 | 10 | 0.0050 | 0.012 | 30 | 0.0310 |
| | 200 | 0.35 | 15 | 0.0030 | 0.014 | 11 | 0.0215 |
| | 300 | 0.35 | 15 | 0.0030 | 0.014 | 74 | 0.0249 |
| VIV2 | **100** | **0.60** | **11** | **0.0045** | **0.010** | **20** | **0.0180** |
| (mode: 49) | 150 | 0.70 | 10 | 0.0050 | 0.012 | 37 | 0.0300 |
| | 200 | 0.60 | 12 | 0.0045 | 0.008 | 39 | 0.0325 |
| | 300 | 0.70 | 11 | 0.0050 | 0.011 | 72 | 0.0440 |

Table S2: Effects of run parameters on performance of DAUMI, UMI-tools, and Calib for simulated data. Underlined numbers indicate the default parameter settings, and the best performance for each method per dataset and metric are bolded, unless there is a tie. For parameters, $d$: threshold of edit distance between UMIs, $e$: error tolerance, $k$: $k$-mer size, $m$: number of minimizer, $t$: minimizer threshold, $\rho$: penalty parameter. The six parameter sets were recommended by Calib for datasets with mean read length 250 and mean barcode length 4 [12]. TP: number of true haplotypes recovered. FP: false positives. Other columns are described in the Table 2.

| **Datasets** | **Method** | **Parameters** | **FP** | **TP** | **b** | **RSS** | $\boldsymbol{R^2}$ |
|---|---|---|---|---|---|---|---|
| Simulation 1 | UMI-tools | $\underline{d = 1}$ | 27 | 22 | **0.931** | **48.450** | **0.995** |
| | UMI-tools | $d = 2$ | **23** | 22 | 0.788 | 67.244 | 0.990 |
| | Calib | $e = 1; k = 8; m = 5; t = 2$ | 29 | 22 | 0.840 | 74.409 | 0.991 |
| | Calib | $e = 1; k = 8; m = 6; t = 2$ | **28** | 22 | 0.846 | 73.409 | 0.991 |
| | Calib | $e = 1; k = 8; m = 6; t = 3$ | **28** | 22 | 0.846 | **66.775** | **0.992** |
| | Calib | $\underline{e = 1; k = 8; m = 7; t = 2}$ | **28** | 22 | 0.843 | 79.077 | 0.990 |
| | Calib | $\underline{e = 1; k = 8; m = 7; t = 3}$ | **28** | 22 | 0.843 | 79.077 | 0.990 |
| | Calib | $e = 1; k = 8; m = 7; t = 4$ | **28** | 22 | **0.850** | 71.957 | 0.991 |
| | DAUMI | $\rho = 1$ | 28 | 22 | 0.994 | 51.657 | 0.995 |
| | DAUMI | $\underline{\rho = 9}$ | 27 | 22 | 0.972 | 43.321 | 0.996 |
| | DAUMI | $\rho = 10$ | 26 | 22 | 0.972 | 42.321 | 0.996 |
| | DAUMI | $\rho = 20$ | 24 | 22 | 0.968 | 41.700 | 0.996 |
| | DAUMI | $\rho = 40$ | **19** | 22 | **0.998** | **40.810** | **0.996** |
| Simulation 2 | UMI-tools | $\underline{d = 1}$ | 21 | 23 | **0.963** | **32.484** | 0.997 |
| | UMI-tools | $d = 2$ | **18** | 23 | 0.935 | 35.550 | 0.997 |
| | Calib | $e = 1; k = 8; m = 5; t = 2$ | 23 | 23 | 0.884 | 71.070 | 0.993 |
| | Calib | $e = 1; k = 8; m = 6; t = 2$ | 23 | 23 | 0.884 | 71.070 | 0.993 |
| | Calib | $e = 1; k = 8; m = 6; t = 3$ | 23 | 23 | 0.884 | 71.070 | 0.993 |
| | Calib | $\underline{e = 1; k = 8; m = 7; t = 2}$ | 23 | 23 | 0.884 | 71.070 | 0.993 |
| | Calib | $\underline{e = 1; k = 8; m = 7; t = 3}$ | 23 | 23 | 0.884 | 71.070 | 0.993 |
| | Calib | $e = 1; k = 8; m = 7; t = 4$ | **22** | 23 | **0.890** | **69.890** | **0.994** |
| | DAUMI | $\rho = 1$ | 28 | 23 | 1.015 | 76.834 | 0.995 |
| | DAUMI | $\rho = 10$ | 28 | 23 | 1.015 | 33.834 | 0.998 |
| | DAUMI | $\rho = 20$ | 27 | 23 | 1.010 | 34.503 | 0.998 |
| | DAUMI | $\rho = 40$ | 17 | 23 | 0.995 | 23.691 | 0.998 |
| | DAUMI | $\underline{\rho = 45}$ | **16** | 23 | **0.995** | **22.691** | **0.998** |
| Simulation 3 | UMI-tools | $\underline{d = 1}$ | 44 | 22 | **0.549** | 135.946 | **0.967** |
| | UMI-tools | $d = 2$ | **42** | 22 | 0.477 | **128.747** | 0.959 |
| | Calib | $e = 1; k = 8; m = 5; t = 2$ | 29 | 22 | 0.680 | 186.068 | 0.970 |
| | Calib | $e = 1; k = 8; m = 6; t = 2$ | 31 | 22 | 0.658 | 187.988 | 0.968 |
| | Calib | $e = 1; k = 8; m = 6; t = 3$ | 30 | 22 | 0.673 | **184.127** | **0.970** |
| | Calib | $\underline{e = 1; k = 8; m = 7; t = 2}$ | **28** | 22 | 0.694 | 327.521 | 0.951 |
| | Calib | $\underline{e = 1; k = 8; m = 7; t = 3}$ | **28** | 22 | 0.694 | 327.521 | 0.951 |
| | Calib | $e = 1; k = 8; m = 7; t = 4$ | 29 | 22 | **0.698** | 324.900 | 0.951 |
| | DAUMI | $\rho = 1$ | 21 | 22 | **0.975** | 61.111 | 0.995 |
| | DAUMI | $\rho = 10$ | 21 | 22 | 0.971 | 39.291 | 0.997 |
| | DAUMI | $\rho = 20$ | 20 | 22 | 0.968 | 43.559 | 0.997 |
| | DAUMI | $\rho = 40$ | 13 | 22 | 0.947 | 29.233 | **0.998** |
| | DAUMI | $\underline{\rho = 49}$ | **10** | 22 | 0.932 | **29.173** | 0.997 |
| Simulation 4 | UMI-tools | $\underline{d = 1}$ | 44 | 21 | **0.651** | **120.385** | **0.964** |
| | UMI-tools | $d = 2$ | **39** | 21 | 0.463 | 134.589 | 0.947 |
| | Calib | $e = 1; k = 8; m = 5; t = 2$ | 42 | 22 | 0.661 | 253.790 | 0.950 |
| | Calib | $e = 1; k = 8; m = 6; t = 2$ | 41 | 22 | 0.643 | 269.999 | 0.945 |
| | Calib | $e = 1; k = 8; m = 6; t = 3$ | **40** | 22 | **0.665** | **243.061** | **0.953** |
| | Calib | $\underline{e = 1; k = 8; m = 7; t = 2}$ | 41 | 22 | 0.651 | 303.096 | 0.940 |
| | Calib | $\underline{e = 1; k = 8; m = 7; t = 3}$ | 41 | 22 | 0.651 | 303.096 | 0.940 |
| | Calib | $e = 1; k = 8; m = 7; t = 4$ | 42 | 22 | 0.654 | 301.254 | 0.940 |
| | DAUMI | $\rho = 1$ | 31 | 22 | **0.972** | **53.410** | **0.995** |
| | DAUMI | $\rho = 10$ | 31 | 22 | 0.936 | 69.267 | 0.993 |
| | DAUMI | $\underline{\rho = 18}$ | 25 | 22 | 0.880 | 96.708 | 0.989 |
| | DAUMI | $\rho = 20$ | 25 | 22 | 0.865 | 107.398 | 0.987 |
| | DAUMI | $\rho = 40$ | **19** | 22 | 0.734 | 83.868 | 0.986 |

Table S3: Performance with known haplotypes $\mathcal{H}$ and varying penalty parameter $\rho$ in simulation. Fitting the linear model, Deduplicated Abundance = $b\times$(True Abundance), we report $R^2$: proportion of variance explained, expected to be 1 for perfect estimation; RSS: residual sum of squares, expected to be 0 for perfect estimation; $b$: coefficient of true abundance, expected to be 1 ($>1$ for overestimation and $<1$ for underestimation); FP: Number of false positives (there are no false negatives); TP: Number of true positives; Dist.: centered scaled distance between $\rho$ and expected amplified abundance, $(\rho -$ Mean)/std.dev. The theoretical Mean and std.dev are reported in Table 1. The best performer for each dataset is bolded unless there is a tie of more than two values of $\rho$.

| Simulation | $\rho$ | Dist. | FP | TP | $b$ | RSS | $R^2$ |
|---|---|---|---|---|---|---|---|
| 1 | 0.01 | -1.74 | 0 | 22 | **0.999** | 14.980 | 0.9987 |
| | 40 | -0.53 | 0 | 22 | 0.988 | **7.277** | **0.9993** |
| 2 | 0.01 | -4.38 | 0 | 23 | 1.016 | 4.426 | 0.9997 |
| | 40 | -2.42 | 0 | 23 | **1.015** | **2.834** | **0.9998** |
| 3 | 0.01 | -4.38 | 2 | 22 | 0.971 | 21.770 | 0.9982 |
| | 10 | -3.89 | 0 | 22 | **0.971** | 16.770 | 0.9986 |
| | 20 | -3.40 | 0 | 22 | 0.969 | 16.443 | **0.9987** |
| | 30 | -2.91 | 0 | 22 | 0.966 | 16.976 | 0.9986 |
| | 40 | -2.42 | 0 | 22 | 0.963 | **15.214** | **0.9987** |
| | 50 | -1.93 | 0 | 22 | 0.946 | 33.594 | 0.9971 |
| 4 | 0.01 | -1.74 | 1 | 22 | **0.972** | **15.466** | **0.9985** |
| | 10 | -1.44 | 0 | 22 | 0.943 | 32.181 | 0.9968 |
| | 20 | -1.13 | 0 | 22 | 0.887 | 76.858 | 0.9913 |
| | 30 | -0.83 | 0 | 22 | 0.830 | 74.992 | 0.9903 |
| | 40 | -0.53 | 0 | 22 | 0.772 | 79.438 | 0.9881 |
| | 50 | -0.23 | 0 | 22 | 0.738 | 101.062 | 0.9836 |

Table S4: Information on the two tested real datasets. Strand, forward or reverse strand used in the analysis; UMI Len. (nt): UMI length in nucleotides; Read Len. (nt), length of sampled sequence in nucleotides; The length of whole read is UMI Len. + Read Len.. No. reads, total number of reads in dataset.

| **Dataset** | **Accession** | **Region** | **Strand** | **UMI Len. (nt)** | **Read Len. (nt)** | **No. reads** |
|---|---|---|---|---|---|---|
| V1V2 | SRR2241783 | V1V2 | Reverse | 9 | 241 | 33.5k |
| V3 | SRR5105420 | V3 | Reverse | 10 | 249 | 530.6k |

Table S5: Agreement of DAUMI results on five random halvings of the V3 dataset as a function of $\rho$. Columns are as described for Table 3. The chosen $\rho$ and the best performance for each column are bolded.

| | | **Deduplicated abundance $\geq$ 1** | | | | **Deduplicated abundance $\geq$ 2** | | | |
|---|---|---|---|---|---|---|---|---|---|
| **Dataset** | $\rho$ | **Hap1** | **Hap2** | **Jaccard** | **Ruzicka** | **Hap1** | **Hap2** | **Jaccard** | **Ruzicka** |
| V1V2 | 1 | 90 (5) | 91 (3) | 0.70 (0.03) | 0.81 (0.02) | 46 (3) | 47 (3) | 0.64 (0.04) | 0.81 (0.02) |
| | **10** | 90 (5) | 90 (4) | 0.72 (0.03) | **0.85** (0.01) | 36 (1) | 37 (3) | 0.77 (0.04) | **0.87** (0.01) |
| | 20 | 88 (4) | 89 (3) | 0.73 (0.04) | 0.84 (0.01) | 36 (1) | 37 (2) | 0.76 (0.03) | 0.86 (0.01) |
| | 40 | 87 (4) | 88 (3) | **0.73** (0.04) | 0.84 (0.01) | 36 (1) | 36 (2) | **0.77** (0.03) | 0.86 (0.01) |
| V3 | 1 | 183 (9) | 174 (4) | 0.39 (0.01) | **0.83** (0.01) | 96 (1) | 96 (4) | 0.74 (0.04) | **0.90** (0.01) |
| | **10** | 180 (9) | 171 (4) | 0.39 (0.01) | 0.72 (0.01) | 68 (4) | 72 (3) | 0.72 (0.03) | 0.83 (0.01) |
| | 20 | 179 (9) | 171 (4) | 0.40 (0.01) | 0.68 (0.02) | 63 (2) | 65 (4) | 0.73 (0.03) | 0.74 (0.02) |
| | 40 | 171 (9) | 164 (2) | **0.40** (0.01) | 0.65 (0.01) | 58 (2) | 59 (3) | **0.75** (0.05) | 0.81 (0.01) |

Table S6: Agreement of Calib, UMItools, Starcode-umi results on five random halvings of V1V2 and V3 datasets after removing singleton UMIs. Columns are as described for Table 3.

(a) Mean (standard deviation)

| | | **Deduplicated abundance $\geq$ 1** | | | | **Deduplicated abundance $\geq$ 2** | | | |
|---|---|---|---|---|---|---|---|---|---|
| **Dataset** | **Method** | **Hap1** | **Hap2** | **Jaccard** | **Ruzicka** | **Hap1** | **Hap2** | **Jaccard** | **Ruzicka** |
| V1V2 | Calib | 183 (6) | 180 (3) | 0.45 (0.02) | 0.69 (0.01) | 37 (1) | 38 (5) | 0.76 (0.04) | 0.86 (0.03) |
| | DAUMI | 90 (5) | 90 (4) | 0.72 (0.03) | **0.84** (0.02) | 37 (1) | 37 (3) | 0.77 (0.05) | 0.87 (0.02) |
| | Naïve | 293 (10) | 303 (7) | 0.29 (0.01) | 0.57 (0.01) | 43 (2) | 43 (3) | 0.75 (0.03) | 0.88 (0.03) |
| | Starcode-umi | 16 (2) | 14 (1) | **0.75** (0.05) | 0.81 (0.02) | 11 (1) | 10 (1) | 0.71 (0.05) | 0.81 (0.02) |
| | UMI-tools | 187 (4) | 199 (2) | 0.50 (0.02) | 0.74 (0.01) | 39 (1) | 40 (3) | **0.83** (0.05) | **0.90** (0.01) |
| V3 | Calib | 1091 (26) | 1094 (24) | 0.05 (0.00) | 0.08 (0.00) | 25 (1) | 21 (2) | 0.40 (0.04) | 0.54 (0.02) |
| | DAUMI | 180 (9) | 171 (4) | **0.39** (0.01) | **0.72** (0.01) | 68 (4) | 72 (3) | **0.72** (0.03) | **0.83** (0.01) |
| | Naïve | 1561 (9) | 1543 (10) | 0.06 (0.00) | 0.22 (0.00) | 91 (4) | 84 (3) | 0.64 (0.04) | 0.74 (0.02) |
| | Starcode-umi | 329 (16) | 321 (9) | 0.06 (0.00) | 0.50 (0.00) | 26 (5) | 26 (3) | 0.14 (0.02) | **0.83** (0.01) |
| | UMI-tools | 1308 (18) | 1324 (18) | 0.05 (0.00) | 0.14 (0.00) | 53 (4) | 51 (4) | 0.70 (0.04) | 0.75 (0.03) |

(b) Change in mean from Table 3

| | | **Deduplicated abundance $\geq$ 1** | | | | **Deduplicated abundance $\geq$ 2** | | | |
|---|---|---|---|---|---|---|---|---|---|
| **Dataset** | **Method** | **ΔHap1** | **ΔHap2** | **ΔJaccard** | **ΔRuzicka** | **ΔHap1** | **ΔHap2** | **ΔJaccard** | **ΔRuzicka** |
| V1V2 | Calib | -625 | -617 | 0.37 | 0.46 | -6 | -5 | 0.09 | 0.04 |
| | DAUMI | | | | | | | | |
| | Naïve | | | | | | | | |
| | Starcode-umi | -2288 | -2297 | 0.75 | 0.62 | -4 | -3 | -0.02 | 0.01 |
| | UMI-tools | -355 | -339 | 0.36 | 0.38 | -3 | -2 | 0.06 | 0.03 |
| V3 | Calib | -3957 | -3927 | 0.05 | -0.03 | -39 | -45 | -0.15 | -0.17 |
| | DAUMI | | | | | | | | |
| | Naïve | | | | | | | | |
| | Starcode-umi | -14748 | -14776 | 0.06 | 0.41 | -70 | -77 | 0.06 | 0.06 |
| | UMI-tools | -2627 | -2400 | 0.05 | 0.00 | -19 | -23 | 0.06 | 0.00 |
